# Fiber-type vulnerability and proteostasis reprogramming in skeletal muscle during pancreatic cancer cachexia

**DOI:** 10.1101/2025.09.15.676415

**Authors:** Bowen Xu, Aniket S. Joshi, Meiricris Tomaz da Silva, Silin Liu, Ashok Kumar

## Abstract

Cachexia is a debilitating syndrome marked by progressive skeletal muscle wasting, commonly affecting cancer patients, particularly those with pancreatic cancer. Despite its clinical significance, the molecular mechanisms underlying cancer cachexia remain poorly understood. In this study, we utilized single-nucleus RNA sequencing (snRNA-seq) and bulk RNA-seq, complemented by biochemical and histological analyses, to investigate molecular alterations in the skeletal muscle of the KPC mouse model of pancreatic cancer cachexia. Our findings demonstrate that KPC tumor growth induces myofiber-specific changes in the expression of genes involved in proteolytic pathways, mitochondrial biogenesis, and angiogenesis. Notably, tumor progression enhances the activity of specific transcription factors that regulate the mTORC1 signaling pathway, along with genes involved in translational initiation and ribosome biogenesis. Skeletal muscle-specific, inducible inhibition of mTORC1 activity further exacerbates muscle loss in tumor-bearing mice, highlighting its protective role in maintaining muscle mass. Additionally, we uncovered novel intercellular signaling networks within the skeletal muscle microenvironment during pancreatic cancer-induced cachexia. Together, these results reveal previously unrecognized molecular mechanisms that regulate skeletal muscle homeostasis and identify potential therapeutic targets for the treatment of pancreatic cancer-associated cachexia.

## Introduction

Cancer cachexia is a multifactorial syndrome characterized by a significant loss of skeletal muscle mass, often accompanied by adipose tissue wasting. It contributes to reduced responsiveness to antineoplastic therapies, diminished quality of life, and lower overall survival in cancer patients (1, 2). While cachexia is common in individuals with gastroesophageal, lung, and head and neck cancers, it is particularly prevalent in pancreatic ductal adenocarcinoma (PDAC) where it is present in approximately 60% of patients at diagnosis and up to 85% in advanced stages (1, 3–6). In pancreatic cancer, cachexia arises from a combination of systemic inflammation, metabolic dysregulation, pancreatic exocrine and endocrine dysfunction, and the activity of hormones, neuropeptides, and tumor-derived catabolic factors (7). Despite its high prevalence and clinical impact, there are currently no effective therapies approved for the prevention or treatment of cancer cachexia, partly because etiology of cancer-induced cachexia remains less understood (5).

Disruption of proteostasis is one of the important mechanisms leading to skeletal muscle wasting in many catabolic conditions, including cancer (4, 8). Many proinflammatory cytokines and tumor-derived factors increase muscle proteolysis through stimulating ubiquitin-proteasome system (UPS) and autophagy (9). In addition to increased proteolysis, other pathological changes also occur in skeletal muscle during cancer cachexia. For example, mitochondrial dysfunction resulting in reduced oxidative phosphorylation and heightened oxidative stress are common pathological features of skeletal muscle in animal models and patients with cachectic cancer (1, 10). Moreover, cancer cachexia involves disruption of neuromuscular junctions leading to functional denervation and the loss of skeletal muscle mass and function (2, 11). It has been reported that the loss of BMP signaling and up-regulation of myogenin are some of the important mechanisms that contribute to the NMJ dysfunction and muscle wasting during cancer cachexia (11, 12). Furthermore, a recent study demonstrated endothelium dysfunction and loss of skeletal muscle vascular density in multiple animal models and in patients with cancer cachexia (13).

Although it is generally believed that the rate of protein synthesis is reduced in skeletal muscle during cancer progression, some studies have shown that protein synthesis remains unaffected, or even increases in animal models and patients with cancer cachexia (14). In mammalian cells, translation is initiated by the binding of eukaryotic initiation factor 4E (eIF4E) to the 5′ cap of the mRNA, followed by the recruitment of eIF4G, eIF4A, and eIF4B. This assembly forms the eIF4F complex, also known as the translation preinitiation complex (15). The mTORC1-p70S6K is a major signaling mechanism that stimulates translation initiation in mammalian cells, including skeletal muscle in response to many growth factors and physical exercise (16–19). Indeed, mTORC1 is required for mechanical load-induced myofiber hypertrophy (20). While mTORC1-mediated signaling is not required for the maintenance of muscle mass, it plays an important role in regulating mitochondrial proteome and muscle contractile force production in response to Akt-mediated myofiber hypertrophy (21). Intriguingly, recent studies employing genetic mouse models have provided strong evidence that constitutive activation of mTORC1 results in muscle wasting and myopathy without having any significant effect on protein synthesis (22, 23). However, the role of mTORC1-mediated signaling in the regulation of muscle mass during pancreatic cancer cachexia remains completely unknown.

While myofibers are the major cell type that are present in skeletal muscle, there are many other cell types, including fibroblasts, endothelial cells, and inflammatory immune cells which may also be affected and contribute to the loss of muscle mass during cancer cachexia (24). Because of their relative low abundance, until recently it was difficult to understand the molecular changes that occur in these other cell types in skeletal muscle during various pathological conditions, including cancer. However, with the advent of single-cell and single nucleus RNA-seq (snRNA-Seq) technologies, it is now possible to study the changes in the transcriptome that occur in individual cell types within skeletal muscle and decipher potential intercellular communications that may be important for the regulation of skeletal muscle mass. Indeed, recent studies have used the snRNA-Seq approach to decipher the role of a few molecular pathways that may regulate muscle mass during cancer cachexia (12, 24). Furthermore, snRNA-Seq has been used to understand the transcriptional changes that occur in skeletal muscle of a mouse model of Lewis lung carcinoma (25). However, the molecular changes that regulate proteostasis and intercellular communications in skeletal muscle during pancreatic cancer cachexia remain completely unknown.

A recent transcriptomic study has identified several molecular and signaling alterations in the skeletal muscle of male pancreatic cancer patients especially at the late stage of the disease that closely resemble those observed in KRAS^G120^P53^R172H^Pdx-Cre^+/+^ (KPC) tumor-bearing mice, a well-established model of pancreatic cancer-associated cachexia (26–28). In the present study, we employed snRNA-seq and bulk RNA-seq, along with various biochemical and histological techniques, to investigate cell type-specific alterations in the skeletal muscle of KPC tumor-bearing mice. Our results demonstrate that KPC tumor growth causes fiber-type reprograming and distinctly affects the activation of specific catabolic and anabolic signaling and proteostasis in skeletal muscle of adult mice. The results also demonstrate that mTORC1 signaling, translational initiation, and ribosome biogenesis are significantly upregulated in fast-type myofibers of KPC tumor-bearing mice. Targeted inhibition of mTORC1 signaling through deletion of Raptor in adult mice exacerbates muscle wasting in response to tumor growth. Finally, snRNA-seq transcriptome analysis has identified novel intercellular communications that may play important roles in muscle wasting during pancreatic cancer cachexia.

## RESULTS

### Single-nucleus RNA-Seq reveals cellular heterogeneity in skeletal muscle during pancreatic cancer cachexia

To investigate the transcriptional changes occurring in skeletal muscle during pancreatic cancer cachexia, we employed the orthotopic mouse model of KPC pancreatic cancer (26). Briefly, 2 x 10^5^ KPC cells were injected into the tail of the pancreas of 12-week-old wild-type mice. Control mice were injected with PBS into the pancreas. On day 21 after KPC cells injection, the mice were analyzed for changes in body weight and grip strength followed by collection of hindlimb muscles and performing histological, biochemical, or transcriptome analyses **(Fig. 1A)**. Relative to day 0, there was a significant reduction in the absolute body weight (BW) and tumor-free BW of KPC tumor-bearing mice compared to control mice **(Fig. 1B, C)**. In addition, there was a significant decrease in the four-paw grip strength normalized by tumor-free BW of KPC tumor-bearing mice compared to control mice **(Fig. 1D)**. Next, transverse sections of TA muscle were generated and immunostained for laminin protein followed by measurement of myofiber cross-sectional area (CSA). Results showed that the average myofiber CSA was significantly reduced in TA muscle of KPC tumor-bearing mice compared with control mice **(Fig. 1E, F)**. We then pooled gastrocnemius (GA) muscle from 5 control and 5 KPC tumor-bearing mice and performed snRNA-seq. High-quality nuclei from PBS (8,953) and KPC (9,794) groups were selected for downstream analyses following the inclusion criteria of nuclei expressing more than 200 but less than 8000 genes, less than 50% mitochondrial gene content, and less than 5% expression of hemoglobin-related genes. Unsupervised clustering of the integrated dataset resulted in 21 spatially distributed clusters which were assigned to 13 distinct cell type identities based on the expression pattern of specific marker genes **(Fig. S1A)**, as previously described (12, 25, 29). For instance, the unique expression of *Pax7* for muscle stem cells (‘MuSCs’), *Myh7* for type I myofibers (‘Type I’), *Myh2* for type IIA myofibers (‘Type IIa’), *Myh4* for type IIB myofibers (‘Type IIb’), *Myh1* for Type IIX myofibers (‘Type IIx’), *Pecam1* for endothelial cells (‘EC’), *Pdgfra* for fibroblasts (‘Fibroblasts’), *Ptprc* for immune cells (‘Immune’), *Col22a1* for myotendinous junctions (‘MTJs’), *Chrne* for neuromuscular junctions (‘NMJs’), *Cidec* for adipocytes (‘Adipocytes’), *Mpz* for schwann cells (‘Schwann’), and *Myh11* for smooth muscle cells (‘Smooth’) was confirmed for accurate annotation of cell-types (**Fig. 1G, H**). Interestingly, our analysis showed the presence of nuclear clusters (i.e., cluster 2, henceforth Cachectic-1; and cluster 14, henceforth Cachectic-2) that were exclusively present in the KPC group **(Fig. 1I)**. Since the gene expression of skeletal muscle-specific E3 ubiquitin ligases, MAFbx (gene name: *Fbxo32*) and MuRF1 (gene name: *Trim63*) were markedly enriched in these clusters, we annotated the cellular identity as ‘cachectic’ nuclei clusters **(Fig. 1G, H** and **Fig. S1A)**. Similar to cachectic myonuclei clusters, there were more than one cluster for Type IIb (i.e., clusters 0, 1, 3, and 9), Fibroblasts (i.e., clusters 4 and 16), and EC (i.e., clusters 7, 10, 12, and 18) nuclei **(Fig. S1A)**. To further investigate the multi-clustering of myonuclei, specifically in cachectic and Type IIb clusters, we characterized functional deviations by identifying differentially expressed genes across nuclei of corresponding cell type-clusters. This analysis showed that the enriched genes in cachectic-1 and 2 nuclear clusters shared association with common pathways such as KEAP1-NFE2L2, positive regulation of catabolic processes, and proteolysis involved in protein catabolic process, indicating that the transcriptome of the two cachectic nuclear clusters was largely similar. By contrast, we observed a few distinctly associated processes with the enriched genes in cachectic-1 and -2 clusters. For instance, protein processing in the ER and regulation of TOR signaling were enriched terms associated with cachectic-1 cluster, whereas cellular response to nutrient levels and glycogen metabolism were associated with cachectic-2 cluster **(Fig. S1B)**. Similarly, we observed that the enriched genes in four clusters of Type IIb (i.e., Type IIb1-4) myonuclei were associated with mitochondrial respiration/oxidative phosphorylation (OXPHOS), supramolecular fiber organization, proteolysis/ER-stress response, and hormone-mediated signaling respectively **(Fig. S1C)**, suggesting functional variance among type IIb myonuclei. While type IIb myofibers are known to be predominantly glycolytic in nature, we observed that OXPHOS-related genes were enriched in a small fraction of type IIb myonuclei (type IIb-1 cluster) which were spatially located away from the other three type IIb clusters. Interestingly, using spatial metabolomics approach, a recent study also showed the presence of a sub-type of type IIb myofibers that are enriched in fatty acid oxidative metabolism (30). We also investigated whether there was any gross effect in the cell type distribution by analyzing the proportion of nuclei in each annotated cluster. The cachectic cluster comprised of ∼24% of all nuclei in the KPC group, which was absent from the PBS group. This led to a biased reduction in the proportion of nuclei of all other cell types, except for the MTJs and NMJs clusters **(Fig. 1J)**.

**Figure 1.**
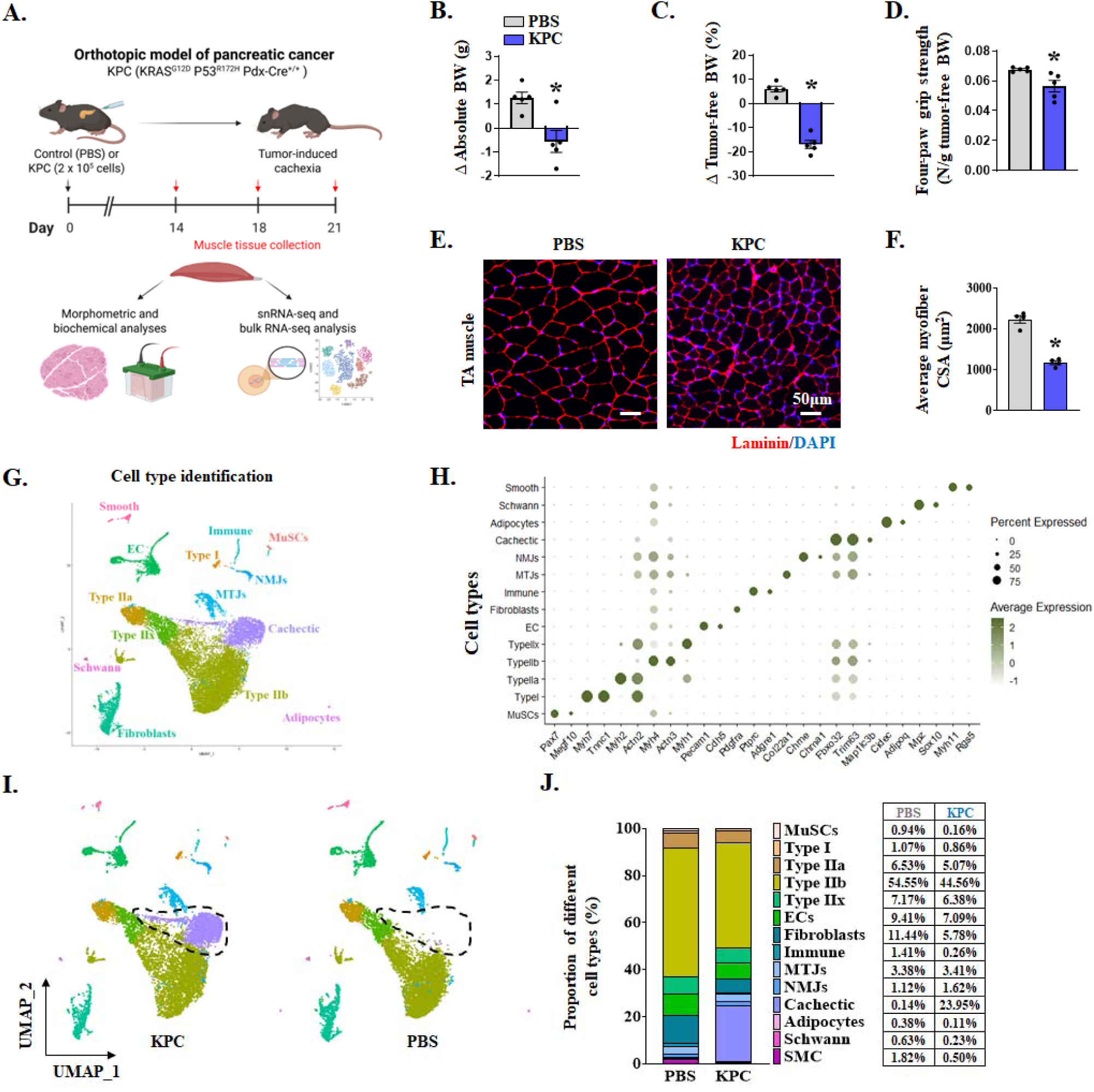
Cellular diversity in skeletal muscle from control and KPC tumor-bearing mice. **(A)** Schematic representation of experimental design. 12-week-old male C57BL6 were injected with KPC cells or PBS into the pancreas. On day 21, hindlimb muscles of the mice were isolated and processed for morphometric, biochemical, and snRNA-Seq analyses. Changes in **(B)** absolute body (BW) weight, and **(C)** tumor-free BW in control and KPC tumor-bearing mice. **(D)** Four-paw grip strength normalized by tumor-free BW of control and KPC tumor-bearing mice. **(E)** Representative anti-laminin and DAPI stained TA muscle transverse sections. Scale bar, 50µm. **(F)** Quantification of average myofiber cross-sectional area (CSA) in TA muscle of control and KPC tumor-bearing mice. n=5 in each group. All data are presented as mean ± SEM. \**p* ≤ 0.05, significantly different from control mice injected with PBS alone analyzed by unpaired Student *t* test. **(G)** UMAP plot representing manually annotated clusters for cell type identity. **(H)** Expression of representative marker genes define major cell types in dot plot. **(I)** Split-UMAP illustrating transcriptomic clustering of muscle-derived nuclei of control and KPC tumor-bearing mice. **(J)** Proportion of different nuclei population in GA muscle of control and KPC tumor-bearing mice.

### Pancreatic cancer triggers atrophy program in skeletal muscle

To characterize the cachectic cluster in KPC tumor-bearing mice, differential gene expression analysis of nuclear clusters of control and KPC tumor-bearing mice was performed. For this analysis, the cachectic cluster in KPC tumor-bearing mice was compared with all myonuclei (i.e., Type I, IIa, IIb, IIx, MTJs, NMJs and MuSCs) of control mice to enrich muscle-specific transcriptional deviation in response to KPC tumor growth. Using the threshold of |Log2FC| > 0.58 and *p*-value < 0.05, our analysis revealed differential regulation of 4360 genes, out of which 1811 genes were upregulated, and 2549 genes were downregulated. Volcano plot shows the distribution of differentially expressed genes (DEGs) in nuclei of cachectic versus all muscle nuclei **(Fig. S2A)**. Gene Ontology (GO) pathway enrichment analysis of DEGs showed that the expression of genes associated with KEAP1-NFE2L2 pathway, protein catabolic process, autophagy, mitophagy, and response to endoplasmic reticulum (ER) stress was highly up-regulated in cachectic nuclei compared to all myonuclei of control mice **(Fig. 2A)**. The Kelch-like ECH-associated protein 1 (Keap1)-nuclear factor erythroid 2-related factor 2 (Nfe2l2 or Nrf2) pathway is a protective mechanism in skeletal muscle that is elicited in response to oxidative stress during cancer cachexia. Indeed, it has been reported that muscle-specific deletion of Nrf2 exacerbates muscle wasting during cancer cachexia (31, 32). Skeletal muscle proteolysis during cancer cachexia is mediated by the ubiquitin-proteasome system (UPS) and autophagy (1, 33, 34). Gene set scoring analysis using the Seurat pipeline revealed that pathway scores for the UPS and autophagy were highly elevated in cachectic and type IIb nuclear clusters, moderately increased in type IIa and IIx clusters, and modestly elevated in type I clusters of KPC tumor-bearing mice compared to control mice (**Fig. 2B and S2B, C**), indicating fiber type-specific upregulation of UPS and autophagy-related gene expression. Consistently, analysis of bulk RNA-Seq dataset revealed that the gene expression of multiple molecules involved in UPS and autophagy/mitophagy is significantly upregulated in skeletal muscle of KPC tumor-bearing mice compared to control mice (35). Furthermore, several genes, including *Fbxo32*, *Trim63*, *Eda2r*, *Il6ra*, and *Ddit4*, whose products are known to promote muscle atrophy (36–39), were among the top 50 upregulated genes in cachectic nuclei compared to all myonuclei of control mice **(Fig. 2C)**. In addition, several of the top 50 upregulated genes in cachectic nuclei were also enriched at the muscle tissue level as illustrated in the bulk RNA-seq heatmap **(Fig. 2D)**. We also performed an independent qPCR analysis of a few mRNAs found to be upregulated in both the snRNA-seq and bulk RNA-seq datasets using muscle samples from control and KPC tumor-bearing mice. Results showed that the mRNA levels of *Sik1*, *Arrdc3*, *Sesn1*, *Retreg1*, *Fbxo32*, and *Trim63* were significantly increased in skeletal muscle of KPC tumor-bearing mice compared to controls **(Fig. 2E)**. Western blot analysis showed that the levels of ubiquitin (Ub)-conjugated proteins, MAFbx, MuRF1, and LC3B-II, but not Beclin1, were significantly increased in TA muscle of KPC tumor-bearing mice compared with control mice **(Fig. 2F and Fig. S2D)**. Together, our transcriptomic analysis identified several new molecules that are highly upregulated in cachectic myonuclei along with the components of UPS and autophagy in response to KPC tumor growth.

**Figure 2.**
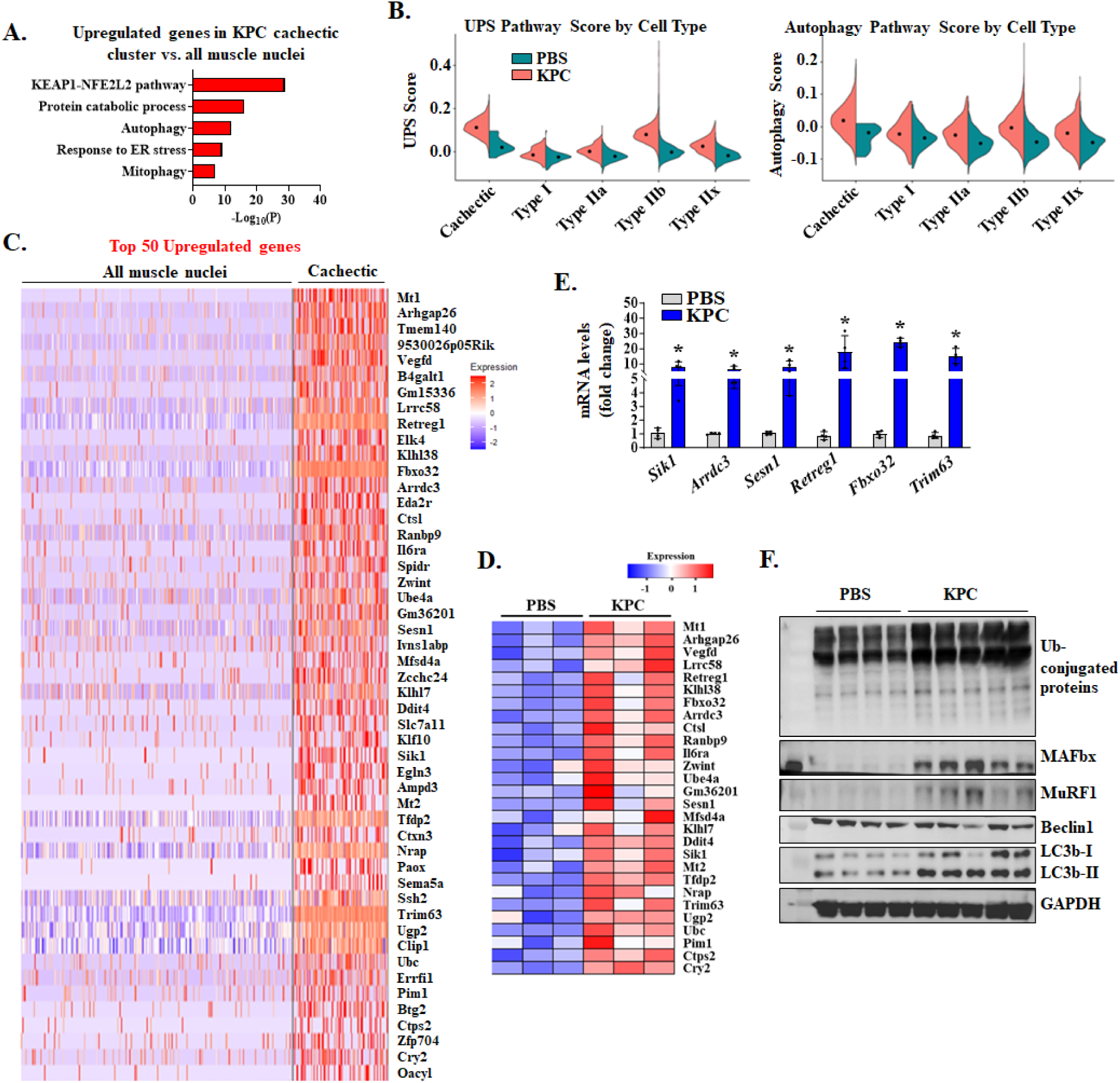
Upregulation of proteolytic systems in cachectic myofibers. **(A)** Enrichment analysis of upregulated genes in cachectic myonuclei highlighting significantly enriched pathways related to protein degradation. **(B)** Violin plots showing module scores for the ubiquitin-proteasome system (UPS) and autophagy across different myonuclear clusters from GA muscle of control and KPC tumor-bearing mice. Scores were calculated using Seurat’s scoring function to quantify pathway activity. Each violin represents the distribution of pathway scores within a fiber type. **(C)** Heatmap displaying the expression of the top 50 upregulated genes in cachectic nuclei compared to all the myonuclei in control mice. (D) Heatmap showing the expression of representative top 50 upregulated genes from Bulk RNA-seq at tissue level. **(E)** Independent qPCR analysis of the upregulated top-ranked representative genes in GA muscle of control and KPC tumor-bearing mice. **(E)** Immunoblots of the levels of total ubiquitinated proteins, MAFbx, MuRF1, Beclin1, LC3B-I/II and unrelated protein GAPDH in TA muscle of control and KPC tumor bearing mice. n=4 for PBS group and n=5 in KPC tumor group. All data are presented as mean ± SEM. \**p* ≤ 0.05, values significantly different from PBS-injected control mice analyzed by unpaired Student *t* test.

### KPC tumor growth represses mitochondrial metabolism, angiogenesis, and structural maintenance genes in skeletal muscle

We next analyzed the downregulated genes in cachectic nuclear cluster of KPC tumor-bearing mice. GO pathway enrichment analysis revealed that gene expression of many molecules involved in aerobic respiration, cytoskeleton in muscle cells, actin-filament based process, circulatory system process, and tube morphogenesis was significantly reduced in cachectic nuclear cluster compared to all myonuclei of control mice **(Fig. 3A)**. Cancer cachexia involves degradation of select thick filament proteins in skeletal muscle (40). Consistently, analysis of top 50 downregulated genes in cachectic cluster versus all myonuclei of control mice showed that the gene expression of multiple molecules related to muscle structure and muscle contraction, including *Myl1*, *Myh4*, *Myh1*, *Tpm1*, *Mylk4*, *Mylpf*, and *Itgb6*, was significantly downregulated in cachectic myonuclei **(Fig. 3B)**. Furthermore, gene set scoring analysis revealed a significant reduction in the expression of oxidative phosphorylation (OXPHOS)-related genes across cachectic and other myonuclear clusters, irrespective of muscle fiber type in KPC tumor-bearing mice compared to controls. In addition, angiogenesis scores were also reduced in cachectic and all other myonuclear clusters of KPC tumor-bearing mice, apart from the type I myonuclear cluster **(Fig. 3C, Fig. S3A, B)**. We also analyzed bulk RNA-Seq dataset from GA muscle of control and KPC tumor-bearing mice. Heatmaps showed that the gene expression of several molecules involved in OXPHOS, or angiogenesis were significantly downregulated in GA muscle of KPC tumor-bearing mice compared with control mice (**Fig. 3D**). To verify the changes in transcripts associated with the deregulated pathways in cachectic nuclei, we performed independent biochemical and histological analyses of skeletal muscle from control and KPC tumor-bearing mice. Western blot analysis showed that the levels of myosin heavy chain (MyHC) were significantly reduced in skeletal muscle of mice in response to KPC tumor growth **(Fig. 3E, Fig. S3C)**. Furthermore, protein levels of some of the mitochondrial respiratory chain complexes, including complex I (CI), CIV, and CIII, were significantly reduced in skeletal muscle of KPC tumor-bearing mice compared with control mice **(Fig. 3F and Fig. S4C)**. Next, transverse sections of soleus muscle were generated and used for immunostaining for CD31 (a widely used marker for studying vascular density) and laminin (to mark myofiber boundaries) protein. There was a marked reduction in the number of CD31^+^ cells per unit area in skeletal muscle of KPC tumor-bearing mice compared to control mice **(Fig. 3G, H)**. These results suggest that the expression of various genes whose products are involved in OXPHOS, angiogenesis, and myofiber structural and contractile function is diminished in skeletal muscle during pancreatic cancer cachexia.

**Figure 3.**
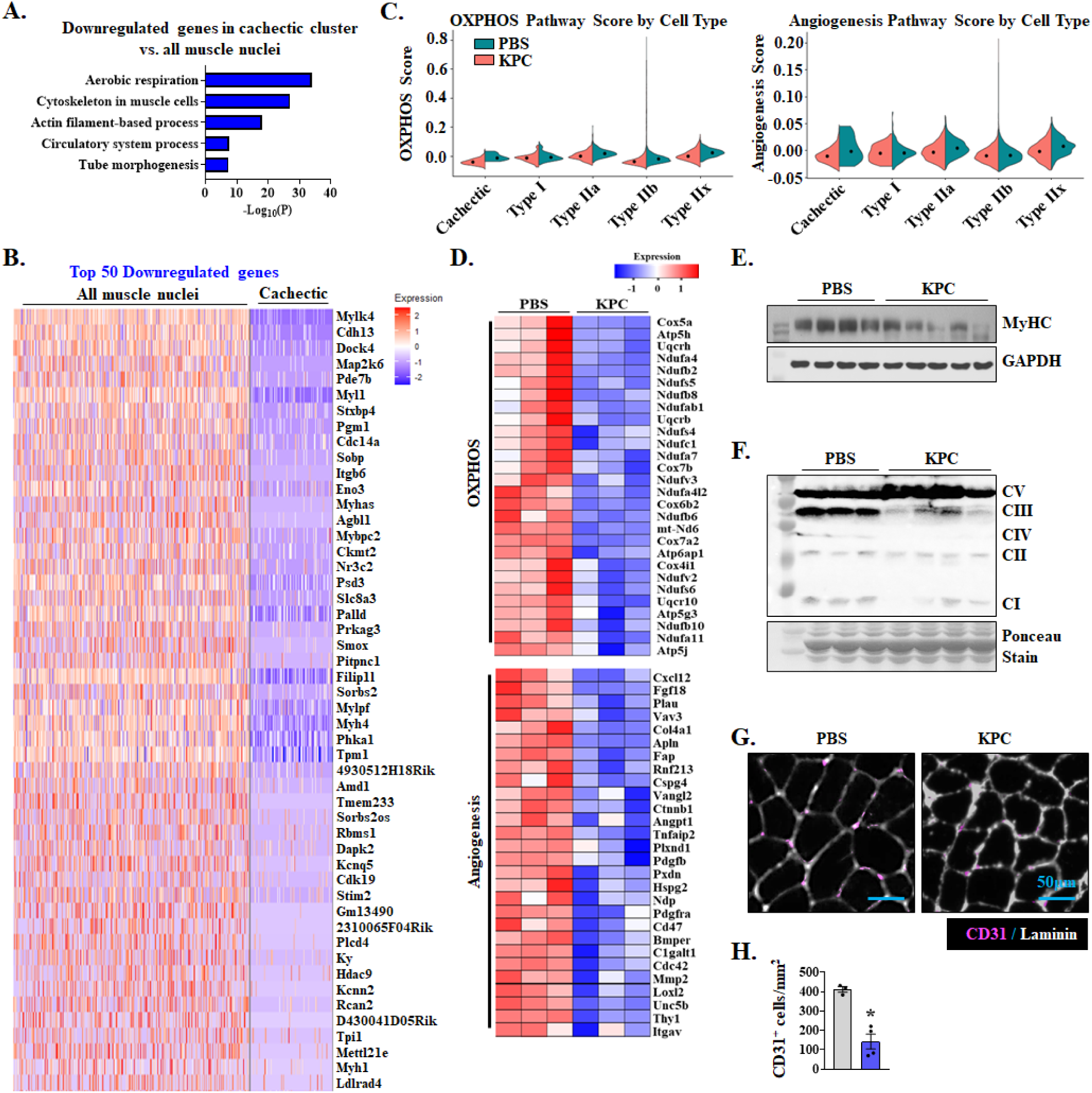
Downregulation of oxidative phosphorylation and angiogenesis pathway in cachectic myofibers. **(A)** Enrichment analysis of downregulated genes in cachectic myonuclei highlighting significantly enriched pathways related to mitochondria and circulation, with top categories including aerobic respiration, cytoskeleton in muscle cells, and circulatory system process. **(B)** Heatmap displaying the expression of the top 50 downregulated genes in cachectic nuclei compared to all the myonuclei of control mice. **(C)** Violin plots showing module scores for the oxidative phosphorylation (OXPHOS) and angiogenesis processes across different myonuclear clusters from GA muscle of control and KPC tumor-bearing mice. Scores were calculated using Seurat’s scoring function to quantify pathway activity. Each violin represents the distribution of pathway scores within a fiber type. **(D)** Heatmap showing the expression of representative genes related to OXPHOS and Angiogenesis from bulk RNA-seq analysis of GA muscle of control and KPC tumor-bearing mice. Immunoblots showing protein levels of **(E)** MyHC, and **(F)** mitochondrial oxidative phosphorylation (OXPHOS) complexes I-V in skeletal muscle of control and KPC tumor-bearing mice. **(G)** Representative photomicrographs (Scale bar, 50 µm) and **(H)** quantification of CD31^+^ cells per unit area after immunostaining for CD31 (pink) and Laminin (white) protein in soleus muscle from control and KPC tumor-bearing mice. n= 3-4 in each group. All data are presented as mean ± SEM. \**p* ≤ 0.05, values significantly different from control mice analyzed by unpaired Student *t* test.

### KPC tumor growth induces fiber-type reprograming in skeletal muscle

Skeletal muscle myofibers are heterogeneous with respect to their expression of MyHC isoform, conferring distinct contractile properties and metabolic function. Based on MyHC isoform expression, it is possible to distinguish type I, IIA, IIX, and IIB fibers (41). While many studies did not find any difference in fiber type distribution, some reported an increase in the proportion of type IIA and IIB myofibers whereas other studies have shown a transition from type IIA towards type IIB in skeletal muscles of cachectic cancer mice (42–45). To determine whether KPC tumor growth alters fiber-type composition and the underlying transcriptional programs in skeletal muscle, we first analyzed the proportion of type I, IIa, IIx, and IIb myonuclei in snRNA-seq dataset. To circumvent the observed bias in the proportion of nuclei **(Fig. 1J)**, we excluded the cachectic clusters and then reanalyzed the proportion of nuclei in all the remaining clusters. This analysis showed that the proportion of type IIx and IIb nuclei was increased in KPC tumor-bearing mice compared with control mice **(Fig. 4A)**. To validate the effect of KPC tumor growth on fiber-type composition, we performed immunohistochemical analysis of various fiber types in TA muscle of control and KPC tumor-bearing mice **(Fig. 4B)**. Results showed that there was a significant reduction in the proportion of type IIa with a concomitant increase in type IIx and IIb myofibers in TA muscle of KPC tumor-bearing mice compared with controls **(Fig. 4C)**. We next investigated how individual muscle fiber-types are affected in response to tumor growth. Using machine learning-based Augur method, nuclei from Type I, IIa, IIx and IIb clusters were ranked based on their sensitivity to cachexia. Results showed that Type IIb myonuclei were the most perturbed, IIx/IIa intermediate, and Type I myonuclei were the least affected by cancer cachexia **(Fig. 4D)**. These results demonstrate that KPC tumor growth induces fiber-type transition of type IIa towards IIb, and that type IIb myofibers are more prone to atrophy.

**Figure 4.**
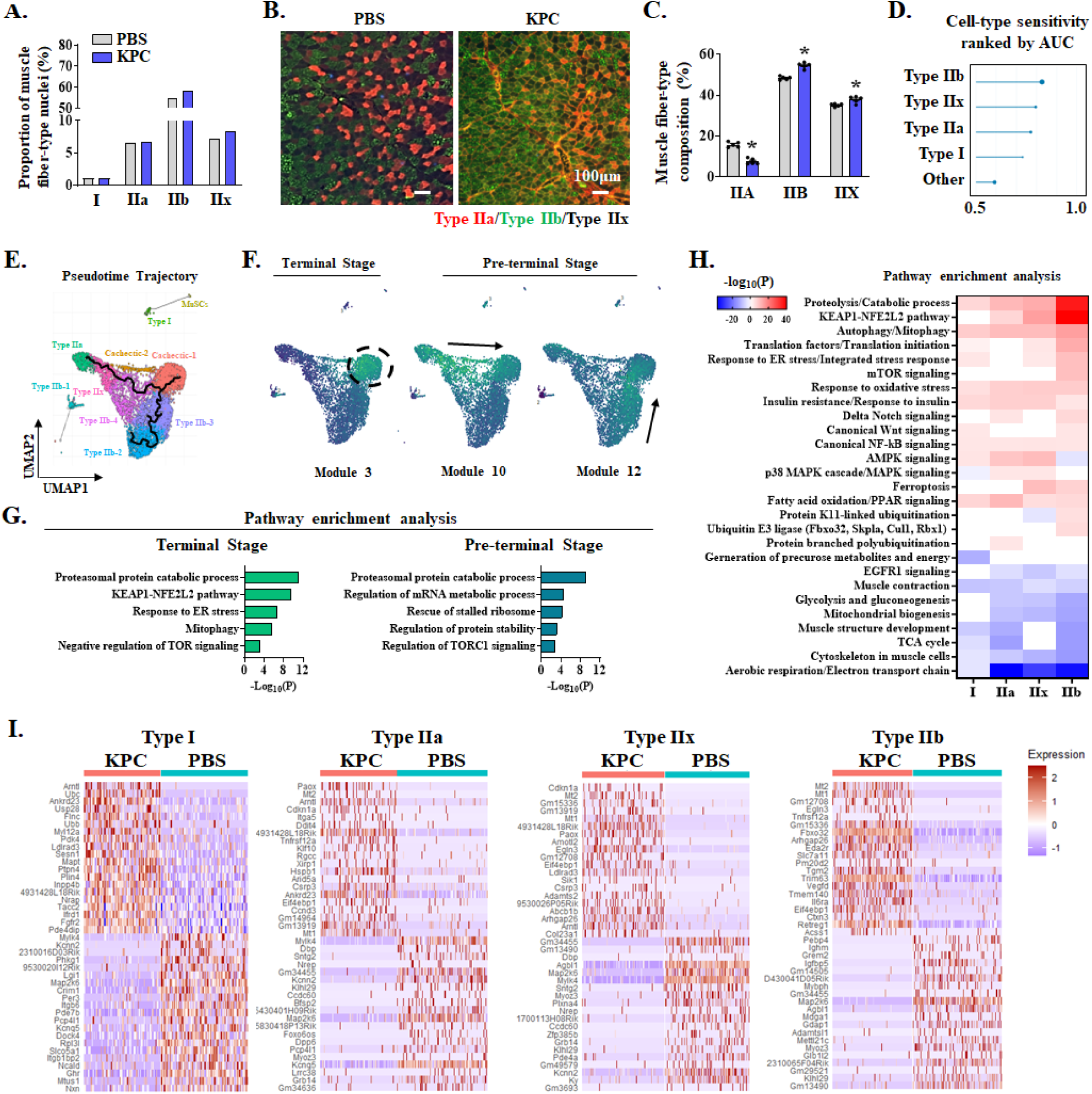
KPC tumor growth causes fiber-type remodeling in skeletal muscle. **(A)** Proportion of muscle fiber type nuclei in snRNA-seq dataset. **(B)** Representative cross-sections of TA muscle from control and KPC tumor–bearing mice after immunostaining for myosin heavy chain (MyHC) isoforms to distinguish fiber types: type IIa fibers (red), type IIb fibers (green), and type IIx fibers (black). Scale bar, 100 µm. **(C)** Quantification of fiber type composition in TA muscle of control and KPC tumor-bearing mice. n=4 in each group. All data are presented as mean ± SEM. \**p* ≤ 0.05, values significantly different from control mice analyzed by unpaired Student t test. **(D)** Augur-based analysis of perturbation sensitivity (AUC, area under the ROC curve) of different fiber-type in skeletal muscle of control and KPC tumor-bearing mice. **(E)** Pseudotime trajectory and annotation of myonuclei subpopulations by UMAP. **(F)** UMAP visualization of module-specific gene expression across myonuclear clusters. Highlighted modules are associated with terminal and pre-terminal fiber stages **(G)** Enrichment analysis of the pathways associated with the pre-terminal/terminal stage gene modules. **(H)** Heatmap showing the pathway enrichment analysis for the DEGs across individual myofiber nuclear cluster. **(I)** The heatmap illustrates the expression of the top 20 upregulated and top 20 downregulated genes in the nuclear cluster of each fiber type.

To further investigate the transcriptional dynamics that regulate the shift of myonuclei towards the cachectic state, we performed trajectory analysis to plot the transition line against a pseudotime axis resembling the development of cachexia. The trajectory line originating from both type IIa and IIb myonuclear clusters reached the terminal node of cachectic cluster. By contrast, type I myonuclear cluster did not exhibit any connection to the cachectic cluster, further suggesting that type I myofibers are less prone to muscle wasting during pancreatic cancer cachexia **(Fig. 4E)**. Since skeletal muscle tissues isolated from KPC tumor-bearing mice are likely to contain myofibers at different stages of atrophy, identification of similarly expressed gene sets/modules in myonuclear clusters can provide deeper insights into the stage-specific progression of cachexia. We then identified co-regulatory gene expression modules in the myonuclear clusters using Louvain community detection analysis. We observed 41 co-regulated gene expression modules in myonuclear clusters (**Fig. S4)**. Interestingly, some of the modules exhibited distinct cachexia-related activity, indicative of differential transcriptional profiles of ‘pre-terminal’ (i.e., approaching cachectic clusters) or ‘terminal’ (i.e., cachectic clusters) stages **(Fig. 4F)**. For instance, Module 3 showed selective enrichment of co-regulated genes within cachectic clusters (terminal stage) which were associated with atrophy program, including proteasome-mediated protein degradation, KEAP1-NFE2L2 signaling, ER-stress responses, mitophagy, and negative regulation of mTOR **(Fig. 4F, G)**. By contrast, enriched genes in Modules 10 and 12 peaked at a pre-terminal divergence zone and were associated with proteasomal protein catabolic process, regulation of mRNA metabolism, ribosome-rescue processes, regulation of protein stability, and regulation of TORC1 signaling, indicative of distinct transcriptional regulation during the progression of cachexia **(Fig. 4F, G)**.

We next investigated how the gene expression of various molecules is regulated in the nuclei of individual muscle fiber-types in control and KPC tumor-bearing mice. Pathway enrichment analysis of DEGs showed that some common top-upregulated pathways in all four types of myonuclear clusters were proteolysis, regulation of catabolic processes, autophagy/mitophagy, fatty acid oxidation/PPAR signaling, response to oxidative stress, insulin resistance, and canonical NF-κB signaling (**Fig. 4H**). Interestingly, some top-upregulated pathways were distinctly regulated in only one or some myonuclear clusters. For example, the expression of genes involved in AMPK signaling was up-regulated in type I, IIa and IIx myonuclear cluster but downregulated in type IIb myonuclei; KEAP1-NFE2L2 pathway in type IIa, IIx, and IIb myonuclear clusters but not in type I cluster. The gene expression of translation factors and the molecules involved in translational initiation was highly elevated in type IIb myonuclei compared to other muscle fiber-types. Moreover, mTOR signaling, protein K11-linked ubiquitination and Ubiquitin E3 ligase were significantly enriched terms observed only in type IIb myonuclei **(Fig. 4H)**. Analysis of downregulated genes showed that a few common pathways, such as those related to aerobic respiration and muscle contraction, were downregulated in all fiber-types. By contrast, the expression of genes involved in mitochondrial biogenesis, glycolysis and gluconeogenesis, and EGFR1 signaling was significantly reduced in all type II myonuclear clusters but not in type I myonuclei **(Fig. 4H)**. The expression of top 20 upregulated and top 20 downregulated genes in each type of myofiber clusters is presented in **Fig. 4I**. Like molecular pathways, there were some common genes that were similarly regulated in more than one myofiber cluster and some others are expressed in distinct clusters. For example, Fn14 (gene name: *Tnfrsf12a*), the receptor for catabolic cytokine TWEAK, is among the top 20 upregulated genes in type IIa and type IIb myonuclear clusters. Indeed, we recently reported that TWEAK-Fn14 system drives muscle wasting during cancer cachexia (46). Similarly, Eda2r, which is also involved in muscle wasting during cancer cachexia (36), is highly upregulated in type IIb myonuclear clusters. Importantly, MAFbx (Fbxo32) and MuRF1 (Trim63) were among the top 20 upregulated genes only in the type IIb myonuclei, further suggesting higher sensitivity of type IIb myofibers in pancreatic cancer cachexia. There were several other molecules, such as Mt1 and Mt2 which are highly expressed in type IIa, IIx, and IIb clusters of KPC tumor-bearing mice, however, their role in the regulation of skeletal muscle mass during cancer cachexia has yet to be investigated **(Fig. 4I)**. Collectively, these data show that KPC tumor growth drives fiber-type transition and distinct transcriptional reprogramming in each type of myofibers in skeletal muscle.

### Transcriptional programs governing protein synthesis are altered in myofibers during pancreatic cancer cachexia

Since transcription factors (TFs) are master regulators of gene expression, we investigated how the activity of different TFs is altered in skeletal muscle of control and KPC tumor-bearing mice. For this analysis, we first identified TF binding motifs overrepresented (i.e., motif enrichment) in the DEGs in myonuclear clusters of our snRNA-Seq dataset. Next, we applied Single-Cell rEgulatory Network Inference and Clustering (SCENIC) which integrates motif enrichment analysis with gene expression data to identify transcription factors and their direct target genes, thereby inferring gene regulatory networks (i.e., regulons). Subsequently, regulon activity was computed at single-cell level (AUCell). This analysis identified several TFs with altered activity in response to KPC tumor growth. The heatmap displays the top 15 most affected TFs (increased and decreased activity) across all myonuclear clusters. **(Fig. 5A)**. We next sought to identify the biological processes associated with the target genes of the deregulated TFs. Pathway enrichment analysis, followed by network-based clustering, revealed multiple interconnected functional modules. Interestingly, we observed several terms associated with the regulation of target of rapamycin complex 1 (TORC1) signaling (highlighted in red) alongside a broader set of translation- and ribosome-related processes (highlighted in blue) **(Fig. 5B)**. We then sorted the TFs based on three selection criteria: elevated TF activity inferred from SCENIC and AUCell analysis, increased expression of TF itself in the cachectic myonuclei, and the overlap of TF regulon with upregulated genes in cachectic clusters. In addition, the sorted TFs were filtered again based on their association with the biological processes highlighted in **Fig. 5B**. The activity of some of the filtered TFs (e.g., Foxo1, Clock, Elk4) was sharply restricted to cachectic myonuclei, whereas others (e.g., Zbtb20, Tead1) display broader induction **(Fig. S5)**, suggesting a layered architecture in which TF modules orchestrate both TORC1 signaling and the downstream translation machinery. An alluvial diagram presents the association of sorted TFs (Tfdp2, Mllt10, Mecp2, Zbtb20, Tead1, Zbtb11, Elk4, Clock, Myf6, Arnt, Arntl, Foxo1, Zfhx3, Zfp639, and Mnt) with the indicated GO terms. The ribbon width signifies the overlapping of target genes associated with the GO terms **(Fig. 5C)**. Interestingly, several biological processes, including ribosome biogenesis, translational initiation, and regulation of post-translational protein modification were linked with multiple transcription factors **(Fig. 5C)**. Since TORC1 is a key signaling molecule regulating protein synthesis and ribosome biogenesis (47), our analysis reveals TORC1 signaling as a central hub in the regulation of muscle proteostasis during pancreatic cancer cachexia.

**Figure 5.**
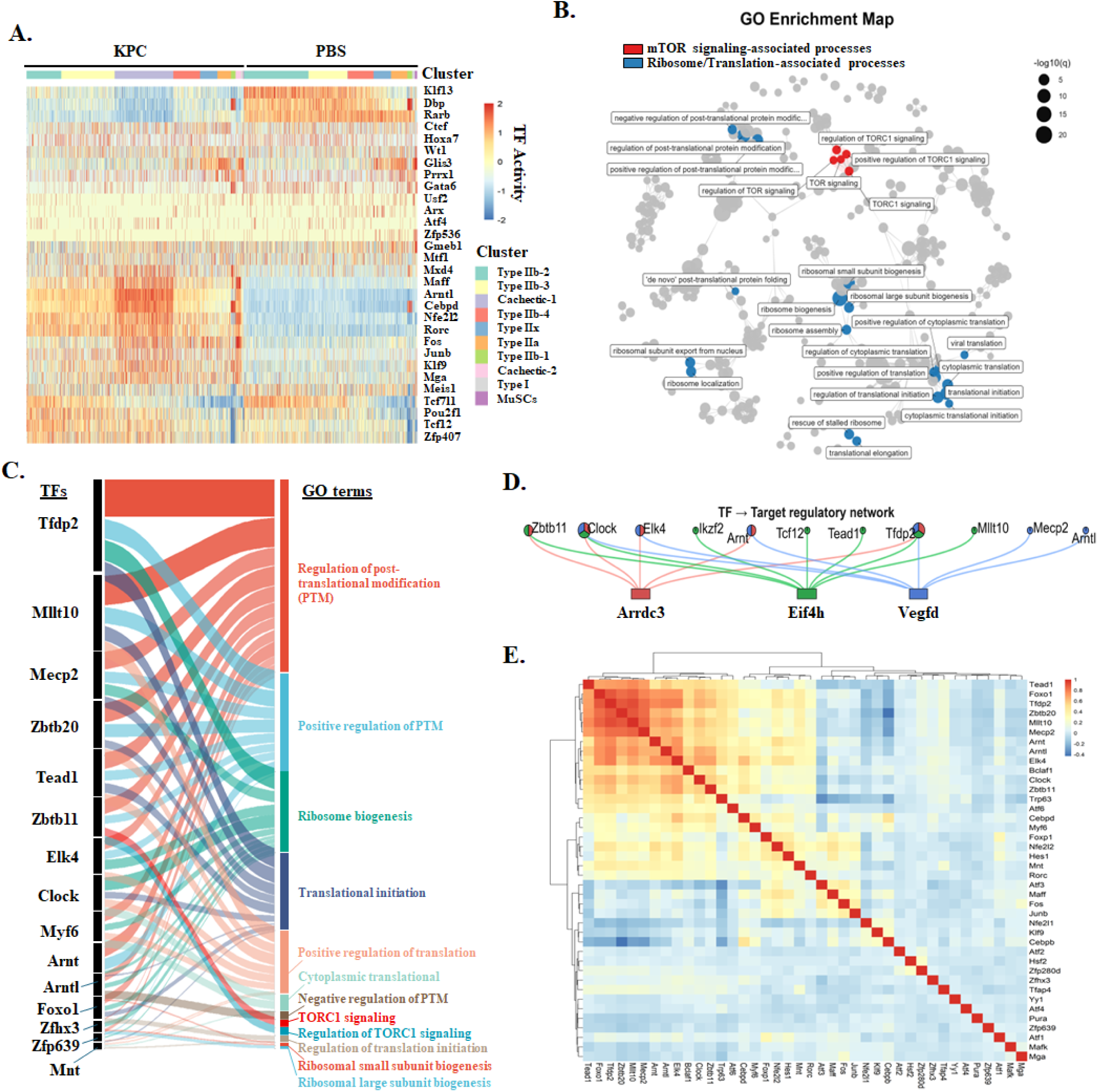
Transcriptional rewiring in cachectic muscle of KPC tumor-bearing mice. **(A)** Heatmap of regulon activity inferred by SCENIC across myonuclei. **(B)** Gene ontology (GO) enrichment network of high-confidence TF targets, with core mTOR signaling terms annotated in red and translation/ribosome-related terms in blue. **(C)** Alluvial diagram linking TFs to enriched GO terms. Ribbon width reflects the number of shared target genes. **(D)** Convergence of enriched TFs regulatory network at key nodes/target genes. **(E)** Correlation matrix of regulon activity scores among deregulated TFs in cachectic myofibers.

We next investigated whether the TFs associated with TORC1-related processes converge on a subset of commonly regulated genes. We observed that distinct TFs converge on 3 target genes, viz. *Arrdc3*, *Eif4h* and *Vegfd* **(Figure 5D)**. Interestingly, Arrdc3 and Vegfd were among the top 50 upregulated genes in cachectic myonuclei compared to myonuclei of control mice **(Fig. 2C)**. Furthermore, eIF4H functionally interacts and stimulates the activity of eIF4A, a component of the eIF4F complex, thereby accelerating protein synthesis (15). Finally, we also investigated pair-wise cooperativity of all the sorted TFs to analyze broad-range correlation in the TF regulatory network. This analysis showed discrete TF cooperative modules in cachectic myofibers. For example, Tead1, Aoxo1, Tfdp2, Zbtb20, Mllt10, Mecp2, and Arnt formed a tightly correlated cluster, indicative of shared or convergent regulatory programs. By contrast, other TFs displayed weak or negative correlations, indicating divergent transcriptional control or potential functional antagonism **(Fig. 5E)**. Taken together, our snRNA-seq analysis identifies novel transcriptional co-regulatory networks and downstream target nodes that impinge on mTOR-related processes in skeletal muscle during pancreatic cancer cachexia.

### Increased translational initiation and ribosome biogenesis in skeletal muscle of KPC tumor-bearing mice

The loss of skeletal muscle mass during cancer cachexia primarily occurs when the rate of protein degradation exceeds the rate of protein synthesis (1, 8). Our analysis of TF activity in cachectic myonuclei suggested that the mTORC1 signaling and protein translation may be affected in skeletal muscle in KPC tumor-bearing mice **(Fig. 5)**. Intriguingly, Gene Set Enrichment Analysis (GSEA) of snRNA-seq dataset revealed that the gene expression of multiple molecules associated with translation initiation and ribosome biogenesis was significantly increased in cachectic myonuclei of KPC tumor-bearing mice compared to all myonuclei of control mice **(Fig. 6A)**. Analysis of both snRNA-Seq and bulk RNA-Seq dataset showed that the gene expression of multiple molecules involved in translation initiation and ribosome biogenesis is significantly upregulated in skeletal muscle of KPC tumor-bearing mice compared to control mice **(Fig. 6B)**. To confirm these findings, we performed independent qPCR and western blot analyses using muscle samples from control and KPC tumor-bearing mice. Results showed that the mRNA levels of various markers of ribosome biogenesis (e.g., 18S rRNA, TIF1a, PAF53, Polr1b, UBF, Rpl5, Rpl11, Rsp3 and Rsp6) were significantly increased in skeletal muscle of KPC tumor-bearing mice compared to control mice **(Fig. 6C)**. By performing surface sensing of translation (SUnSET) assay, we next investigated how the rate of protein synthesis is affected in skeletal muscle in response to KPC tumor growth. Results showed that amounts of puromycin-tagged peptides were significantly increased in skeletal muscle of KPC tumor-bearing mice compared with control mice assayed on day 14 and 18 after implantation of KPC cells **(Fig. 6D and Fig. S6)**. Furthermore, the levels of phosphorylated and total 4EBP1 protein, and phosphorylated p70S6K were also up-regulated in skeletal muscle of KPC tumor-bearing mice compared to control mice further suggesting an increase in translation initiation and protein synthesis in skeletal muscle of KPC tumor-bearing mice. The eIF4H is also one of the three molecules on which various TFs converge in cachectic myonuclei **(Fig. 5D).** Our western blot analysis confirmed that the levels of eIF4H protein are increased in skeletal muscle of KPC tumor-bearing mice **(Fig. 6D and Fig. S6)**.

**Figure 6.**
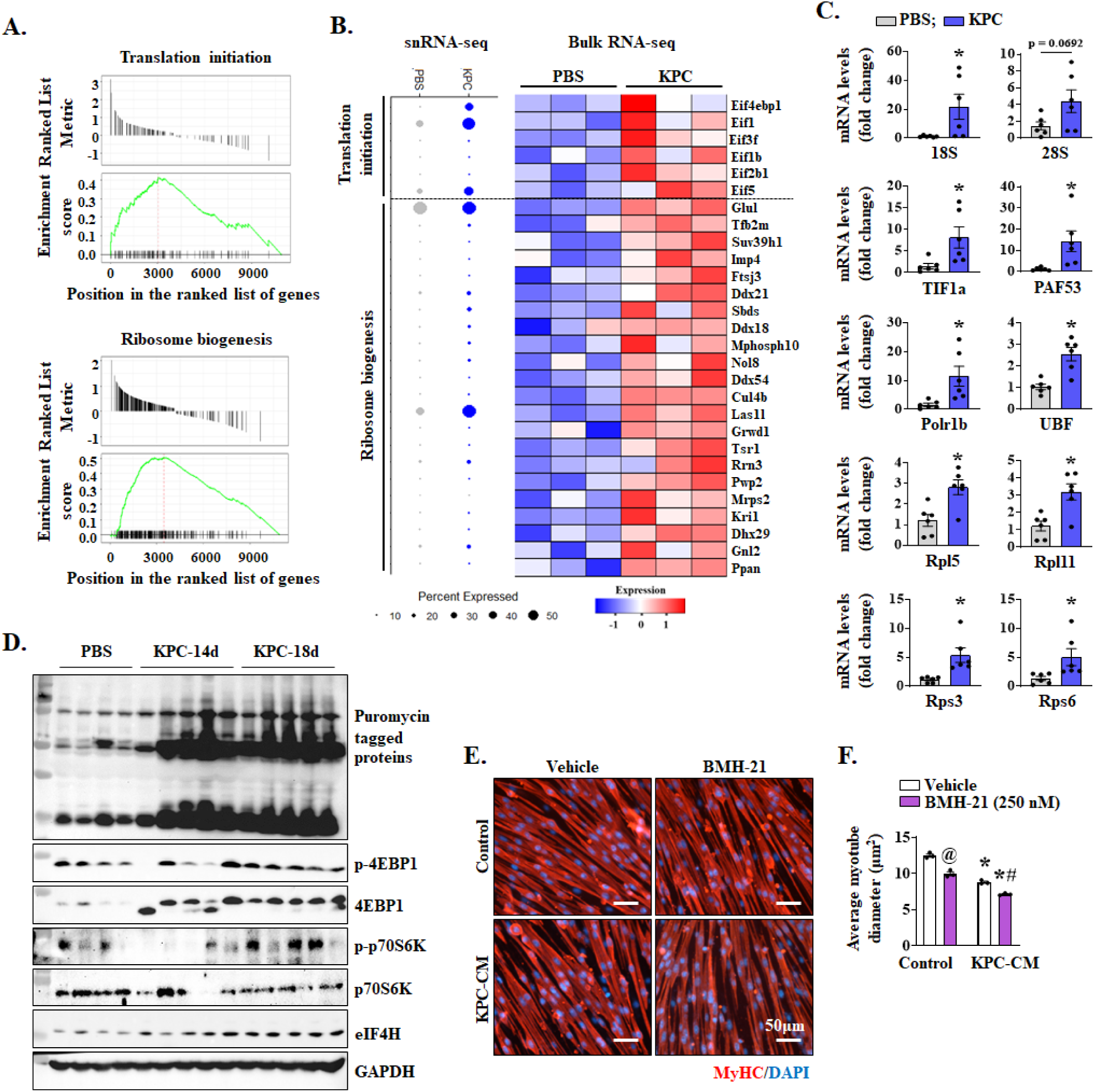
Increased ribosome biogenesis and translation initiation in skeletal muscle of tumor-bearing mice. **(A)** Gene set enrichment analysis (GSEA) showing significant upregulation of translation initiation and ribosome biogenesis processes in cachectic myonuclear clusters compared with all other nuclei population of control mice. **(B)** Dot plots from the snRNA-seq dataset demonstrating the proportion of nuclei (dot size) expressing translation initiation and ribosome biogenesis genes (left) and heatmap showing corresponding expression changes in an independent bulk RNA-seq dataset (right). **(C)** Relative mRNA levels of ribosome biogenesis-related molecules in skeletal muscle of control and KPC tumor-bearing mice. n=4-6 in each group. Data are presented as mean ± SEM. \**p* ≤ 0.05, values significantly different from control mice analyzed by unpaired Student t test. **(D)** Immunoblots showing levels of puromycin-tagged peptides, and levels of p-4EBP1, 4EBP1, p-70S6K, p70S6K, eIF4H protein in TA muscle of control and KPC tumor-bearing mice after 14 and 18 days of inoculation of KPC cells in pancreas. **(E)** Representative photomicrographs and **(F)** quantification of average myotube diameter in cultures treated with vehicle alone or BMH-21 and incubated with or without KPC cells conditioned medium (KPC-CM). n=3 in each group. Data are presented as mean ± SEM. \**p* ≤ 0.05, values significantly different from corresponding control cultures, ^@^*p* ≤ 0.05, values significantly different from vehicle-treated control cultures, and ^#^*p* ≤ 0.05, values significantly different from control cultures incubated in KPC-CM analyzed by two-way ANOVA followed by Tukey’s multiple comparison test.

We have recently reported that KPC cells conditioned media (KPC-CM) induces atrophy in cultured mouse primary myotubes (46). To understand the physiological significance of increased ribosome biogenesis in cachectic muscle, we studied the effect of BMH-21, a small molecule inhibitor of RNA polymerase I that targets the rate-limiting step of ribosome biogenesis (48), on the average diameter of control and KPC-CM-treated cultured myotubes. Briefly, primary myoblasts isolated from mice were incubated in differentiation medium (DM) for 48 h, followed by pre-treatment with vehicle alone or 250 nM BMH-21 for 2 h. Subsequently, the myotube cultures were incubated in exhausted DM (control) or KPC-CM with or without BMH-21 for additional 24 h. The cultures were then fixed and immunostained for MyHC protein followed by measurement of myotube diameter. Results showed that the treatment with BMH-21 alone was sufficient to reduce the average myotube diameter which was further exacerbated when co-incubated with KPC-CM **(Fig. 6E, F)**. Collectively, these results suggest that translational initiation and ribosome biogenesis are enhanced in skeletal muscle during pancreatic cancer cachexia, which may be an adaptive mechanism to restrict the severity of muscle wasting.

### Targeted ablation of Raptor exacerbates muscle wasting during pancreatic cancer cachexia

Since we observed an increase in mTORC1 signaling (**Fig. 5**), we next sought to investigate the effect of inhibition of mTOCR1 signaling on the regulation of muscle mass in response to KPC tumor growth. The regulatory-associated protein of mammalian target of rapamycin (Raptor) binds to mTOR and is essential for mTORC1 activity. Previous studies have shown that muscle-specific deletion of Raptor (gene name: *Rptor*) leads to a drastic reduction in the protein levels and activity of mTORC1 without affecting muscle mass and function under normal physiological conditions (20, 49). Using inducible muscle-specific *Rptor*-knockout mice, we determined whether the inhibition of mTORC1 signaling affects skeletal muscle mass during pancreatic cancer cachexia. Briefly, floxed *Rptor* (henceforth, Rptor^fl/fl^) mice were crossed with HSA-MCM mice to generate Rptor^fl/fl^;HSA-MCM (henceforth, Rptor^mKO^) and littermate Rptor^fl/fl^ mice. 12-week old Rptor^mKO^ mice were treated by i.p. injection of tamoxifen for 4 consecutive days to induce the deletion of Raptor. Littermate Rptor^fl/fl^ mice were also treated with tamoxifen and served as controls. After 2 d of last tamoxifen injection, the mice were injected with PBS alone or 2 × 10^5^ KPC cells into the tail of pancreas. The mice were analyzed on day 21 after inoculation with KPC cells. There was a marked reduction in the wire hanging time and four-paw grip strength normalized by tumor-free body weight in KPC tumor-bearing Rptor^fl/fl^ and Rptor^mKO^ mice compared to corresponding control mice. Although statistical significance was not achieved, there was a trend towards reduced average hanging time and four-paw grip strength in KPC tumor-bearing Rptor^mKO^ compared to KPC tumor-bearing Rptor^fl/fl^ mice **(Fig. 7A, B)**. There was a significant reduction in the wet weight of TA, QUAD, soleus, and GA muscle of both Rptor^fl/fl^ and Rptor^mKO^ mice in response to KPC tumor growth **(Fig. 7C)**. Interestingly, tumor-induced loss in the wet weight of QUAD and GA muscle was significantly higher in Rptor^mKO^ mice compared with Rptor^fl/fl^ mice **(Fig. 7D)**. In contrast, there was no significant difference in tumor wet weight between Rptor^fl/fl^ and Rptor^mKO^ mice **(Fig. S7A)**.

**Figure 7.**
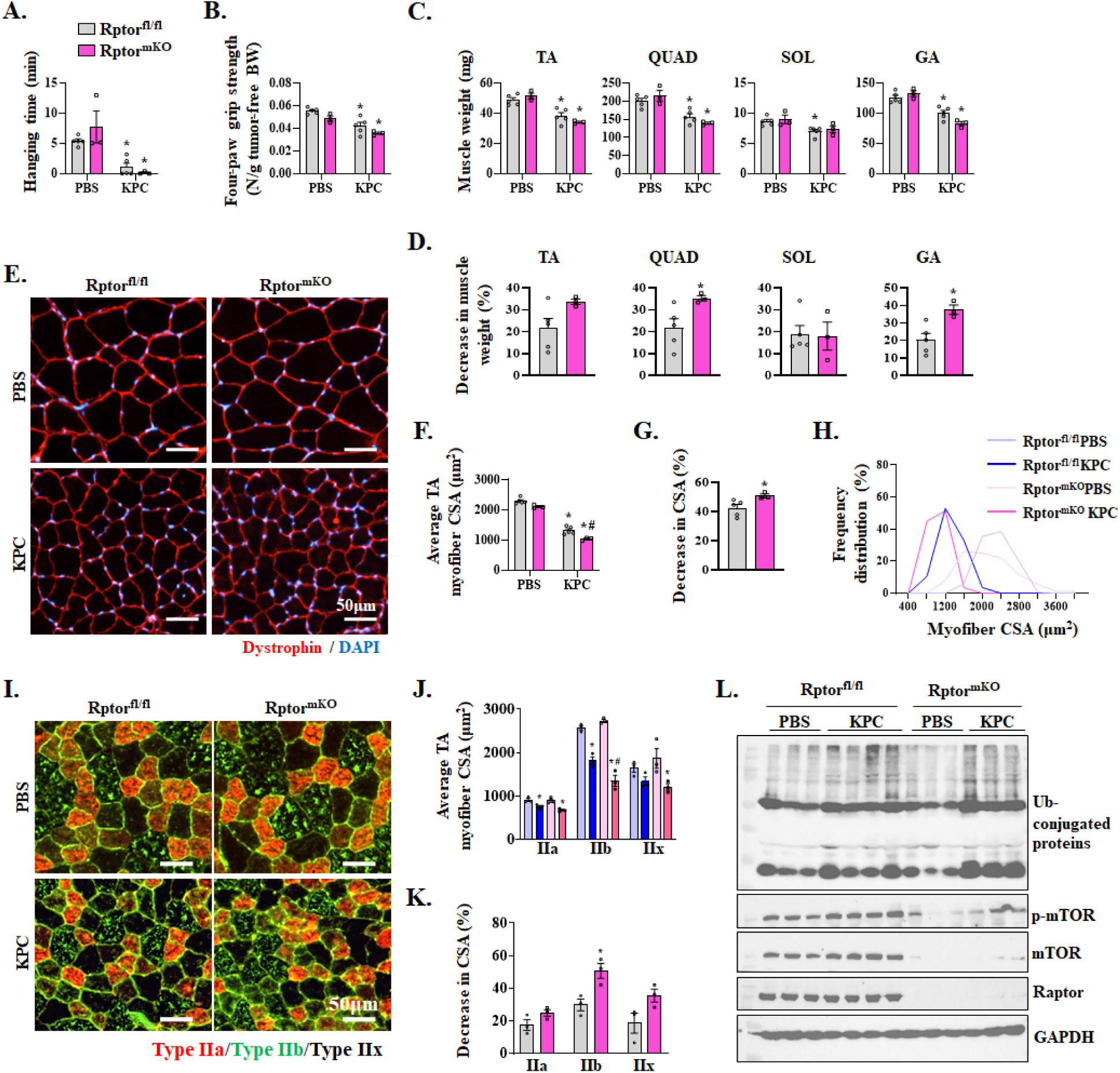
Targeted deletion of Raptor exacerbates muscle loss in response to KPC tumor growth. **(A)** Wire hanging time and **(B)** four-paw grip strength in control and KPC tumor-bearing Rptor^fl/fl^ and Rptor^rmKO^ mice. **(C)** Wet muscle weight of tibialis anterior (TA), quadriceps (QUAD), soleus (SOL), and gastrocnemius (GA) muscle control and KPC tumor-bearing Rptor^fl/fl^ control and Rptor^mko^ mice. **(D)** Decrease in individual muscle weight in Rptor^fl/fl^ and Rptor^mKO^ mice in response to KPC tumor growth. **(E)** Representative images of TA muscle cross-sections after anti-dystrophin (red) and DAPI (blue) staining. Scale bar, 50 µm. **(F)** Quantification of myofiber cross-sectional area (CSA), and **(G)** percentage decrease in average myofiber CSA in TA muscle of Rptor^fl/fl^ and Rptor^mKO^ mice in response to KPC tumor growth. **(H)** Myofiber CSA frequency distribution in TA muscle of control and KPC tumor-bearing Rptor^fl/fl^ and Rptor^mKO^ mice. **(I)** Representative photomicrographs of TA muscle cross-sections after immunostaining for MyHC isoforms: type IIa (red) and type IIb (green). Black/unstained myofibers are type IIx myofibers. Scale bar, 50 µm. **(J)** Quantification of myofiber cross-sectional area (CSA), and **(K)** percentage decrease in average myofiber CSA in individual muscle type of myofibers in TA muscle of Rptor^fl/fl^ and Rptor^mKO^ mice in response to KPC tumor growth. **(L)** Immunoblots showing levels of ubiquitin (Ub)-conjugated proteins, and levels of p-mTOR, mTOR, and Raptor protein in skeletal muscle of control and KPC tumor-bearing Rptor^fl/fl^ and Rptor^mKO^ mice. n=3-5 in each group. All data are presented as mean ± SEM. \**p* ≤ 0.05, values significantly different corresponding control mice, #*p* ≤ 0.05, values significantly different KPC-tumor bearing Rptor^fl/fl^ mice analyzed by unpaired Student *t* test for two-group comparison and two-way ANOVA followed by Tukey’s multiple comparison test for multiple-group data.

Next, transverse sections of TA muscle were generated and used for histological and morphometric analysis by performing H&E staining or immunostaining for dystrophin protein **(Fig. 7E and Fig. S7B)**. There was a significant reduction in the average cross-sectional area (CSA) of both KPC tumor-bearing Rptor^fl/fl^ and Rptor^mKO^ mice compared to corresponding control mice. Importantly, average myofiber CSA of TA muscle of KPC tumor-bearing Rptor^mKO^ mice was significantly less compared to corresponding Rptor^fl/fl^ mice **(Fig. 7F).** Further analysis showed that tumor growth led to a significant reduction in the average myofiber CSA in TA muscle of Rptor^mKO^ mice compared with Rptor^fl/fl^ mice (**Fig. 7G**). Consistently, frequency distribution analysis showed that muscle-specific deletion of Raptor resulted in an apparent increase in the proportion of smaller myofibers in response to KPC tumor growth **(Fig. 7H)**. Since we observed that type IIb myonuclei are more sensitive to cachexia **(Fig. 4D)** and that genes involved in mTOR signaling were upregulated solely in type IIb myonuclei **(Fig. 4H)**, we next investigated whether inhibition in mTORC1 activity in Rptor^mKO^ mice leads to fiber-type-dependent atrophic response in KPC tumor-bearing mice. Immunostaining for different MyHC isoforms showed that the average CSA of all myofiber types (i.e., type IIa, IIb, and IIx) was significantly reduced in TA muscle of both KPC tumor-bearing Rptor^fl/fl^ and Rptor^mKO^ mice compared to corresponding control mice **(Fig. 7I, J)**. Interestingly, KPC tumor-induced reduction in average CSA of type IIb myofibers was significantly higher than that of type IIa and IIx myofibers in Rptor^mKO^ mice compared to Rptor^fl/fl^ mice **(Fig. 7K)**. Deletion of *Rptor* was confirmed by a drastic reduction in the levels of Raptor protein in skeletal muscle of Rptor^mKO^ mice compared to Rptor^fl/fl^ mice. In addition, the levels of both p-mTOR and mTOR protein were also reduced in Rptor^mKO^ mice irrespective of tumor burden. Interestingly, there was also a small increase in the levels of ubiquitin-conjugated proteins in skeletal muscle of Rptor^mKO^ mice compared to Rptor^fl/fl^ mice in response to KPC tumor growth **(Fig. 7L)**. Together, these results suggest that targeted inhibition of mTORC1 exacerbates skeletal muscle wasting during pancreatic cancer cachexia.

### Remodeling of inferred intercellular signaling in skeletal muscle in response to KPC tumor growth

Our previous results focused on the intramuscular transcriptional regulation during pancreatic cancer cachexia. However, in addition to these intrinsic changes, other cell types present in the muscle microenvironment may also contribute to muscle wasting during cancer cachexia. We next investigated the inter-cellular signaling interactions between nuclear clusters identified in our snRNA-seq dataset. For this analysis, we applied CellChat with population weighting to map ligand-receptor mediated intercellular signaling. Results showed that KPC tumor growth induced extensive alterations in both the number and strength of interactions across different muscle and non-muscle cell types in skeletal muscle **(Fig. 8A)**. Further analysis showed that the nuclei in KPC tumor-bearing mice exclusively acquired information flow from various molecules, such as EDA, EDN, PLAU, Nectin, CSPG4, IGFBP, NPR2, TWEAK, ADGRA, WNT, CNTN, CD34 and CD39 which was absent in the nuclear clusters of control mice **(Fig. 8B, Fig. S8)**. In contrast, nuclear clusters of KPC tumor-bearing mice lost information flow from cholesterol, THBS, AGRN, GRN, VCAM, LIFR, SPP1, LAIR1, SEMA5, NPNT, OSTN, VISTA, CD48, CD80, and HH which were present in the control nuclear clusters **(Fig. 8B, Fig. S8)**. We next investigated how these molecules are affected in different nuclei populations of control vs KPC tumor-bearing mice. This analysis showed that there were significant alterations in signaling from different molecules (e.g., IGF1, Visfatin, MSTN) between nuclear clusters of different cell types in control and KPC tumor-bearing mice **(Fig. 8B, Fig. S9A)**. Moreover, CD39 signaling was present in EC and smooth muscle nuclei whereas CD34 was present in only EC of tumor-bearing mice. CNTN and Wnt were present in almost all types of nuclei of tumor-bearing mice, but not in control mice. TWEAK, a potent muscle wasting cytokine (46), was present in cachectic, MTJ, smooth muscle, type IIa and Type IIb nuclear clusters **(Fig. S8C and Fig. S9B)**. We also found that strength of information flow from several other molecules implicated in muscle wasting, such as EDA, Prostaglandin, CD46, NGL, Notch, Testosterone, CCL, MSTN, Visfatin, fibronectin, ANGPT, TGFβ, FLRT, BMP, and FGF was considerably increased in cachectic nuclei of KPC tumor bearing mice even though some of these signals were present in type II myofibers of both control and tumor-bearing mice **(Fig. 8C)**. We performed further analyses to identify the ligand/receptor enrichment in our snRNA-Seq dataset. In control muscle, enrichment was dominated by Col4a3, Mpz, Lama2, Tnxb, and Col4a1, consistent with a structural ECM and neuromuscular junction (NMJ) supportive environment. In contrast, KPC muscle shifted toward Fn1, Col4a1, Col4a2, and Vegfa reflecting ECM remodeling and altered vascular-endocrine communication **(Fig. 8D)**. Together, these integrated metrics demonstrate reprogramming of intercellular signaling in a directional and pathway-specific manner, supporting catabolic alterations such as ECM remodeling, inflammation, and neuromuscular degeneration, in skeletal muscle during pancreatic cancer cachexia.

**Figure 8.**
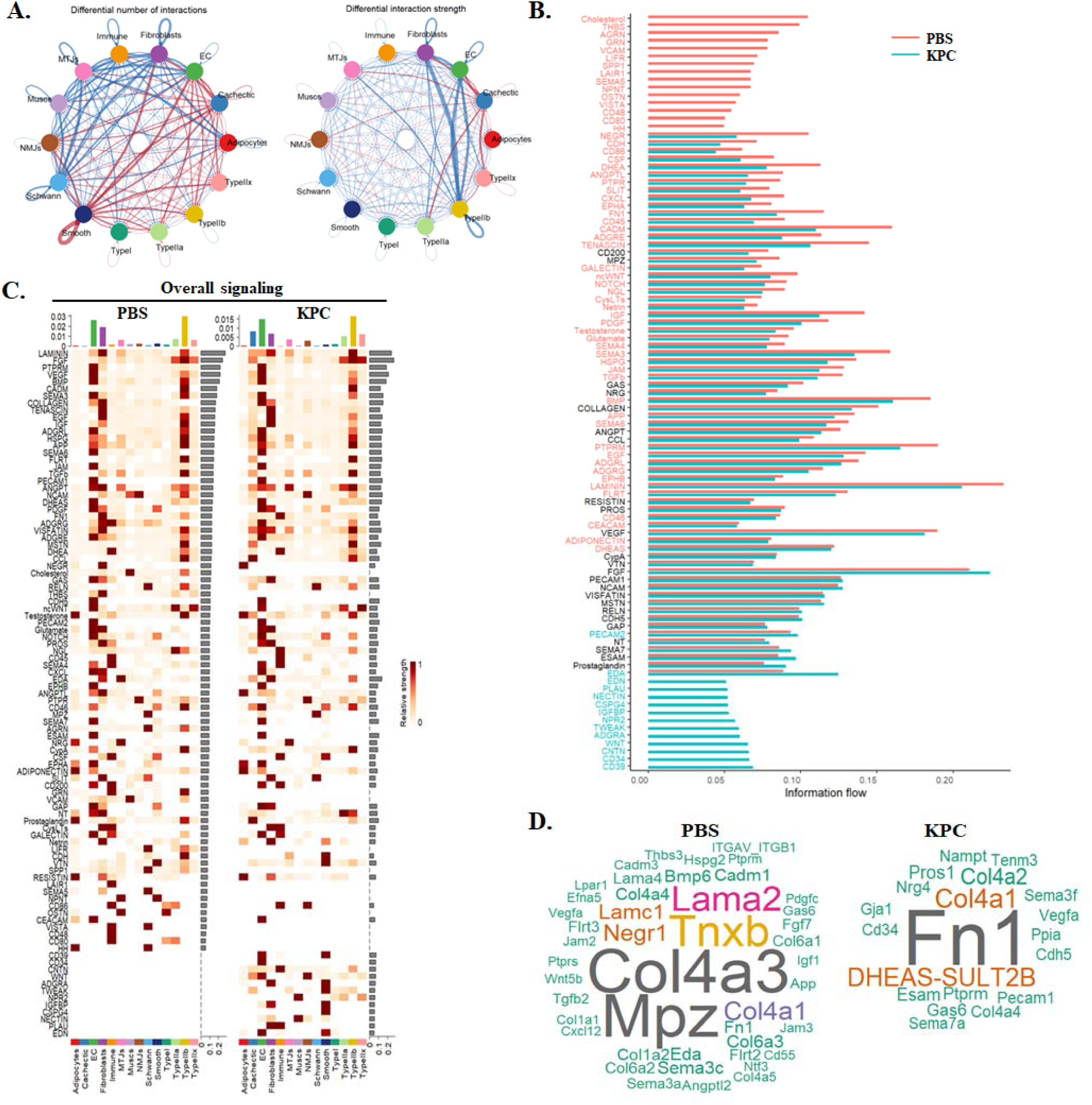
Alterations in inferred intercellular communications in skeletal muscle of KPC tumor-bearing mice. **(A)** Circle plots showing differential number of cellular interactions and strength of interactions between different nuclear clusters of control and KPC tumor-bearing mice. **(B)** Overall information flow comparison of multiple pathways in nuclei of control and KPC tumor-bearing mice. **(C)** Heatmaps of overall signaling (population-weighted) in nuclear clusters of control and KPC tumor-bearing mice. **(D)** Word cloud visualization of ligand/receptor activity highlights dominant receptor or ligand in control and KPC tumor-bearing mice.

## Discussion

Cachexia is major catabolic condition during malignant tumor progression involving the loss of skeletal muscle mass with or without concomitant fat loss (9). Although significant efforts have been made to understand the etiology of cancer cachexia, no drug has been approved so far to treat or prevent this devastating syndrome. Some pharmacological compounds targeting specific pathways have shown promise in improving muscle mass and body weight in animal models. However, clinical trials with these compounds have yielded unsatisfactory results, highlighting the need for a deeper understanding of the underlying molecular mechanisms to develop effective therapies for cancer cachexia (1, 2, 7). In the present study, we have used snRNA-Seq and bulk RNA-Seq approaches to investigate the molecular changes that occur in different cell types present in skeletal muscle microenvironment in the KPC mouse model of pancreatic cancer cachexia. We have identified a unique myonuclei population that we termed as cachectic cluster which expresses the molecular signature of muscle atrophy. Our analysis also demonstrates that the transcriptional regulation, specifically in the type II myofibers, induces transition of myonuclei towards the cachectic state. Furthermore, our results demonstrate that several common and distinct transcriptional changes occur in each myofiber types in response to tumor growth **(Fig. 4)**.

UPS and autophagy are the major mechanisms driving muscle proteolysis in various catabolic conditions, including cancer-induced cachexia (4, 8, 50). Some studies suggest that the type IIb myofibers undergo atrophy at faster rate compared to other types of myofibers in various atrophying conditions, including cancer cachexia (33, 51). Our analysis of the snRNA-Seq dataset showed that molecular signatures of both UPS and autophagy are highly up-regulated in cachectic myonuclei as well as type IIb and IIx myonuclear clusters, suggesting that glycolytic myofibers are more susceptible to wasting compared to oxidative myofibers during cancer cachexia **(Fig. 2, 4)**. Our analysis further showed that the gene expression of many other molecules (e.g., Mt1/Mt2, Vegfd, Elk4, Retreg1, Arrdc3 etc.) is highly induced in cachectic myonuclei. Future studies will characterize the role of these proteins in the etiology of muscle wasting during cancer cachexia.

Mitochondrial dysfunction is a well-established and common pathological feature in the skeletal muscle during cancer cachexia (2, 4, 52). Tumor-derived factors and the host’s inflammatory response trigger several overlapping mechanisms that inhibit mitochondrial biogenesis and cause mitochondrial damage and impaired energy metabolism (52). Damaged mitochondria can also cause oxidative stress which can independently activate catabolic pathways in skeletal muscle (53, 54). Analysis of snRNA-seq dataset revealed that the gene expression of many molecules related to oxidative phosphorylation is diminished in cachectic myonuclei compared to controls **(Fig. 3)**. Recently, it was also reported that the vascular supply is diminished in skeletal muscle during cancer induced cachexia (13). The angiogenic program in skeletal muscle is governed not only by the endothelial and vascular smooth muscle cells, but also by the factors produced by myofibers (55). Furthermore, muscle vasculature is critical for the regeneration of skeletal muscle in response to chronic muscle injury and myofiber hypertrophy following resistance exercise (56–59). The results of the present study demonstrate that the gene expression of several molecules that support angiogenesis is diminished in all myonuclear clusters except in type I myonuclei. Bulk RNA-Seq and histological analysis further confirmed that gene expression of various angiogenesis-related molecules and vascular supply in diminished in skeletal muscle in response KPC tumor growth **(Fig. 3)**.

Skeletal muscle wasting during cancer cachexia involves coordinated activation of a number of signaling pathways and transcription factors which regulate the gene expression of muscle structural proteins and various components of proteolytic systems, mitochondrial biogenesis, and vasculature (1, 4). Our results demonstrate that the activity of specific transcription factors and the expression of their target genes is significantly up-regulated in the cachectic myonuclear clusters of KPC tumor-bearing mice compared with myonuclei of control mice. While the role of some of these TFs (e.g., Foxo1) in the regulation of muscle mass has been elucidated previously, we identified several other TFs which are highly up-regulated and play a co-regulatory role in cachectic myonuclei but their specific role in cancer-induced muscle wasting has not yet been investigated **(Fig. 5)**. Intriguingly, our analysis showed that mTORC1 signaling, ribosome biogenesis, and translation initiation are some of the major processes associated with the activated TFs in cachectic myonuclei of KPC tumor-bearing mice **(Fig. 5B, C)**. A previous study has shown that ribosome biogenesis and protein synthesis are diminished in skeletal muscle of a mouse model of ovarian cancer-induced cachexia (60). In addition, there are other reports which suggest significant reduction, no change, or upregulation of protein synthesis in skeletal muscle of animal models of cancer cachexia (14, 45). Consistent with the snRNA-Seq analysis, our independent biochemical analysis showed that the markers of ribosome biogenesis, translation initiation, and protein synthesis are significantly upregulated in skeletal muscle of KPC tumor-bearing mice **(Fig. 6)**. While the upregulation of protein synthesis and ribosome biogenesis may represent a compensatory response to counteract excessive proteolysis within myofibers, it is not sufficient to prevent muscle atrophy during pancreatic cancer cachexia.

Overexpression of IGF1 or constitutively active Akt is known to induce myofiber hypertrophy and prevent muscle atrophy under various catabolic conditions (61, 62). Akt promotes muscle growth primarily through the activation of the raptor-associated, rapamycin-sensitive mTORC1 complex. Activated mTORC1 phosphorylates 4E-BP1, resulting in its dissociation from the translation initiation factor eIF4E. This allows eIF4E to associate with eIF4G, facilitating the assembly of the eIF4F complex and promoting cap-dependent translation initiation (63, 64). In parallel, mTORC1 phosphorylates p70S6K, which enhances protein synthesis by modulating the activity of multiple components involved in translation. Intriguingly, a few recent studies have demonstrated that the inhibition of mTORC1 through inducible deletion of raptor in skeletal muscle does not prevent the increase in protein synthesis that occurs in response to mechanical overload suggesting that mTORC1-independent mechanisms may also contribute to protein synthesis (20, 65). Furthermore, recent studies have also shown that constitutive activation of mTORC1 for a prolonged period can cause skeletal muscle atrophy and myopathies (22, 23, 66–68). However, the results of the present study demonstrate that the mTORC1-mediated signaling prevents the excessive loss of muscle mass during pancreatic cancer cachexia supported by the findings that targeted inducible deletion of Raptor exacerbates muscle wasting in the KPC tumor bearing mice **(Fig. 7)**. In addition, mTORC1-mediated adaptive response was observed selectively in type IIb myofibers **(Fig. 4H)**, corroborated by the findings that inactivation of mTORC1 results in a profound decrease in average CSA of type IIb myofibers compared with other muscle fiber-types in response to KPC tumor growth **(Fig. 7K)**. These findings are also supported by a previously published report which demonstrates that forced activation of Akt-mTORC1 signaling inhibits skeletal muscle wasting in other mouse models of cancer cachexia (69). Altogether, the results of the present study suggest that skeletal muscle undergoes transcriptional reprograming to enhance translation capacity and efficiency in an attempt to prevent extreme muscle wasting during pancreatic cancer cachexia.

In addition to myofibers, skeletal muscle contains many other cell types which may also play important roles in the regulation of muscle mass in tumor-bearing hosts. Our snRNA-seq analysis demonstrates that tumor growth affects the transcriptome of other cell types, such as endothelial cells, smooth muscle, and fibroblasts present in skeletal muscle microenvironment. Moreover, cell-cell communication is greatly altered in skeletal muscle of tumor-bearing mice compared with controls **(Fig. 8)**. While the signaling from IGF1 is diminished, there is a dramatic increase in the signaling from catabolic molecules, such as TWEAK, EDA, and myostatin in cachectic myonuclei. Interestingly, some of the catabolic molecules are produced predominantly by non-muscle cells which then act on myofibers which express cell surface receptors for them **(Fig. S8, 9)**. This selective rewiring in muscle microenvironment may explain fiber-type vulnerability and establishes a cachectic microenvironment that promotes progressive myofiber loss.

While our study has provided important insights into the molecular mechanisms underlying muscle wasting during pancreatic cancer cachexia, it also has a few limitations. Notably, the snRNA-seq analysis was conducted at a single time point in both control and tumor-bearing mice. Future studies examining gene expression dynamics across different stages of tumor progression would offer a more comprehensive understanding of disease development. Additionally, it remains to be determined whether similar transcriptional changes also occur in skeletal muscle across other models of cancer cachexia. Furthermore, key findings from the snRNA-seq analysis, particularly the involvement of specific transcription factors, warrant further investigation using genetic mouse models of cancer cachexia. Despite these limitations, our study identifies novel mechanisms and potential molecular targets relevant to pancreatic cancer cachexia.

## Methods

### Animals

C57BL/6 wild-type (WT) mice were purchased from Jackson Laboratories (Bar Harbor, ME, USA). Floxed Raptor (*Rptor^fl/fl^*; Strain: B6.Cg-Rptor^tm1.1Dmsa^/J) mice were crossed with HSA-MCM (Strain: B6.Cg-Tg(ACTA1-cre/Esr1*)2Kesr/J, Jackson Laboratory, Bar Harbor, ME) mice to generate muscle-specific Raptor knockout (*Rptor^mKO^*) and littermate control (*Rptor1^fl/fl^*) mice. All mice were in the C57BL/6 background, and their genotype was determined by PCR from tail DNA. We employed KPC orthotopic mouse model of pancreatic cancer cachexia (26, 70). KPC cells, as described (70), were kindly provided by Elizabeth Jaffee (Johns Hopkins University, Baltimore, MD). Briefly, KPC cells (2 × 10^5^ cells in 20 µl PBS) were injected into the tail of the pancreas of 12-week-old WT or *Rptor^fl/fl^* and *Rptor^mKO^* mice. Control mice received an injection of PBS alone. The mice were weighed weekly and euthanized on day 14-21 after the injection of KPC cells. All the animals were handled according to the approved institutional animal care and use committee (IACUC) protocol (PROTO201900043) of the University of Houston. All surgeries were performed under anesthesia, and every effort was made to minimize suffering.

### Cell culture and immunostaining

KPC cells were kindly provided by Dr. Elizabeth Jaffee (Johns Hopkins University, Baltimore, MD) and cultured in RPMI-1640 medium supplemented with 10% fetal bovine serum (FBS). To prepare KPC cell-conditioned medium (KPC-CM), cells were grown to confluency, then incubated for 24 hours in differentiation medium (DM; DMEM supplemented with 2% horse serum). The supernatant was collected, clarified by centrifugation, and filtered through a sterile 0.22 μm syringe filter. For myotube atrophy experiments, KPC-CM was diluted 1:4 in fresh DM.

Primary myoblasts were isolated from hindlimb muscle of C57BL/6 mice as described (71). For myotube formation, primary myoblasts were incubated in DM for 48 h. Myotubes were treated with vehicle alone (DMSO) or BMH-21 for 2h followed by addition of DM (control) or KPC-CM with or without BMH-21 and incubated for additional 24 h. For immunostaining, the cultures were fixed with 4% PFA in PBS for 15 min at room temperature and permeabilized with 0.1% Triton X-100 in PBS for 10 min. Cells were blocked with 2% bovine serum albumin in PBS and incubated with mouse-anti-MyHC (clone MF20, DSHB, Iowa City, Iowa) overnight at 4 °C. The cells were then washed with PBS and incubated with a secondary antibody at room temperature for 1 h. Nuclei were counterstained with DAPI for 3 min. Average myotube diameter was calculated by measuring diameter of 100-120 MyHC^+^-myotubes per group using the NIH ImageJ software. For consistency, diameters were measured at the midpoint along with the length of the MyHC^+^ myotubes.

### Grip strength test

Four-paw grip strength in mice was measured using a digital grip strength meter (Columbus Instruments, Columbus, OH), as previously described (72). Briefly, mice were acclimated for 5 minutes prior to testing. Each mouse was allowed to grasp a metal pull bar with all four paws, after which the tail was gently pulled backward in a horizontal plane until the animal released its grip. The peak force at the moment of release was recorded as the grip strength. Each mouse underwent five trials with 30-second intervals between tests. The average peak force across the five trials was calculated and normalized to body weight.

### Histology and morphometric analysis

Individual TA and soleus muscles were isolated from mice, snap-frozen in liquid nitrogen, and sectioned with a microtome cryostat. For the assessment of muscle morphology, 8-μm-thick transverse sections of TA muscle were stained with hematoxylin and eosin (H&E) dye. Muscle sections were also processed for immunostaining for laminin protein to mark the boundaries of myofibers. Briefly, frozen transverse sections of TA or soleus muscle were fixed in either acetone or 4% paraformaldehyde (PFA) in PBS, followed by blocking with 1% bovine serum albumin (BSA) in PBS for 1 h at room temperature. Sections were incubated overnight at 4°C under humidified conditions with primary antibodies against Type I, Type IIa, and Type IIb myosin heavy chains (1:100; DSHB, University of Iowa, Iowa City, IA), laminin (1:500; Sigma Chemical Co.), or laminin and CD31 protein diluted in blocking solution. After brief washes in PBS, sections were incubated with Alexa Fluor 350-, 488-, or 568-conjugated secondary antibodies (1:500; Invitrogen) for 1 h at room temperature, followed by three 5-min washes in PBS. Nuclei were counterstained with DAPI. Slides were mounted with fluorescence mounting medium (Vector Laboratories) and imaged at room temperature using a Nikon Eclipse Ti-2E inverted microscope equipped with a Digital Sight DS-Fi3 camera and Nikon NIS-Elements AR software. Image levels were uniformly adjusted in Adobe Photoshop CS6 (Adobe). For morphometric analysis, myofiber cross-sectional area (CSA) was analyzed in anti-laminin-stained muscle sections using ImageJ software (NIH, Bethesda, MD). For each muscle, the CSA distribution was determined by analyzing approximately 200 myofibers. Similarly, we quantified the number of CD31 cells per unit area in soleus muscle section using ImageJ software.

### Western blot

Skeletal muscle of mice was washed with PBS and homogenized in lysis buffer (50 mM Tris-Cl (pH 8.0), 200 mM NaCl, 50 mM NaF, 1 mM dithiothreitol, 1 mM sodium orthovanadate, 0.3% IGEPAL, and protease inhibitors). Approximately, 100μg protein was resolved on each lane on 10-12% SDS-PAGE gel, transferred onto a nitrocellulose membrane, and probed using a specific primary antibody (described in **Table S1**). Bound antibodies were detected by secondary antibodies conjugated to horseradish peroxidase (Cell Signaling Technology). Signal detection was performed by an enhanced chemiluminescence detection reagent (Bio-Rad). Approximate molecular masses were determined by comparison with the migration of prestained protein standards (Bio-Rad). All uncropped images are provided in Supplemental **Fig. S9**.

### RNA extraction and qRT-PCR

RNA isolation and qRT-PCR were performed following a standard protocol as described (73). In brief, total RNA was extracted from skeletal muscle of mice using TRIzol reagent (Thermo Fisher Scientific) and RNeasy Mini Kit (Qiagen, Valencia, CA, USA) according to the manufacturer’s protocols. First-strand cDNA for PCR analysis was made with a commercially available kit (iScript cDNA Synthesis Kit, Bio-Rad Laboratories). The quantification of mRNA expression was performed using the SYBR Green dye (Bio-Rad SsoAdvanced-Universal SYBR Green Supermix) method on a sequence detection system (CFX384 Touch Real-Time PCR Detection System, Bio-Rad Laboratories). The sequence of the primers is described in supplemental **Table S2**. Data normalization was accomplished with the endogenous control (β-actin), and the normalized values were subjected to a 2-ΔΔCt formula to calculate the fold change between control and experimental groups.

### Bulk RNA-seq analysis

Total RNA from GA muscle of control and KPC tumor-bearing mice was extracted using TRIzol reagent (Thermo Fisher Scientific) using the RNeasy Mini Kit (Qiagen, Valencia, CA, USA) according to the manufacturer’s protocols. The mRNA-seq library was prepared using poly (A)-tailed enriched mRNA at the UT Cancer Genomics Center using the KAPA mRNA HyperPrep Kit protocol (KK8581, Roche, Holding AG, Switzerland) and KAPA Unique Dual-indexed Adapter kit (KK8727, Roche). The Illumina NextSeq550 was used to produce 75 base paired-end mRNA-seq data at an average read depth of ∼38 M reads/sample. RNA-seq fastq data were processed using CLC Genomics Workbench 20 (Qiagen). Illumina sequencing adapters were trimmed, and reads were aligned to the mouse reference genome Refseq GRCm39.105 from the Biomedical Genomics Analysis Plugin 20.0.1 (Qiagen). Normalization of RNA-seq data was performed using a trimmed mean of M values. Genes with Log2FC ≥ 0.25 and p-value < 0.05 were assigned as differentially expressed genes (DEGs) and represented in volcano plot using ggplot function in R software (v 4.2.2). Pathway enrichment analysis was performed using the Metascape Gene Annotation and Analysis tool (metascape.org). Heatmaps were generated by using heatmap.2 function using z-scores calculated based on transcripts per million (TPM) values. Genes involved in specific pathways were manually selected for heatmap expression plots. All the raw data files can be found on the NCBI SRA repository using the accession code PRJNA1255232.

### snRNA-seq

Freshly isolated GA muscle tissues were processed for the isolation of nuclei. The nuclei were pooled from 5 mice in each group. snRNA-seq libraries were generated using ChromiumTM Single Cell 3’ v3 kit (10X Genomics) according to the manufacturers protocol. Sequencing was performed on an Illumina NovaSeqXPlus system. The sample demultiplexing was performed based on 10 bp i5 and i7 sample index reads to generate Read1 and Read2 paired-end reads. The Cell Ranger Software Suite (10X Genomics, v7.0.0) was used for data demultiplexing, transcriptome alignment, and UMI counting. Raw data are stored in FASTQ (fq) format files, which contain sequences of reads and corresponding base quality. The mouse genome (GRCm38), version M23 (Ensembl 98) was used as reference for read alignments and gene counting with cellranger count. The Seurat R package was further utilized to achieve data quality control.

### snRNA-seq downstream analysis

Cell calling was performed using the EmptyDrops algorithm, which identifies cell-associated barcodes based on unique molecular identifier (UMI) counts and RNA content. Quality control was conducted using Seurat, and nuclei were excluded if they met any of the following criteria: fewer than 200 genes detected, genes with non-zero counts in fewer than 3 nuclei, more than 8000 detected genes (to exclude potential multiplets), mitochondrial gene content greater than 50%, or hemoglobin-related gene content greater than 5%. After filtering, gene expression matrices were normalized by total UMI counts per nucleus, scaled by a sample-specific median UMI factor, and log-transformed using the log1p method. Feature selection was performed to remove uninformative genes and identify biologically relevant signals. Highly variable genes (HVGs) were identified using the FindVariableGenes function in Seurat with the vst method, and the top 3000 HVGs were selected for each sample. Principal component analysis (PCA) was conducted using these HVGs to reduce dimensionality and capture major axes of transcriptomic variation. The resulting principal components were used for unsupervised clustering and identification of cell subpopulations. For visualization, dimensionality was further reduced to two dimensions using t-distributed stochastic neighbor embedding (t-SNE) and uniform manifold approximation and projection (UMAP).

### Gene Module Analysis

Pseudotemporal dynamics were reconstructed using Monocle3 (v1.3.7) (74). Seurat objects from KPC and PBS samples were converted to Monocle3 cell_data_set objects via as.cell_data_set(). Only myonuclei clusters (e.g., Type IIa, IIb, IIx, and cachectic myonuclei) were retained based on prior annotations. Dimensionality reduction was performed using 50 principal components (num_dim = 50), and UMAP coordinates from Seurat were directly transferred to ensure consistency in embedding across analyses. Cells were clustered in Monocle3, and trajectories were learned using the learn_graph() function. Graph-based differential expression testing was performed using graph_test() with the principal graph or kNN graph as the neighborhood structure (neighbor_graph = “principal_graph” or “knn”). Genes with FDR-adjusted q-values < 0.05 were retained. Gene modules were identified via find_gene_modules() using resolution parameters ranging from 10 to 10 ¹ to capture modules at multiple scales. Module expression was aggregated across cell partitions using aggregate_gene_expression(), and module–partition expression heatmaps were visualized using Ward’s method for clustering.

### Comparative Intercellular Communication Analysis

We profiled condition-resolved signaling with CellChat (v2.1.2) as described (75) on a Seurat object split by group labels (PBS, KPC). Cell types were curated by renaming Seurat clusters and stored as meta.data$new_clusters (e.g., Type IIb, Type IIx, Type IIa, Type I, cachectic myonuclei, fibroblasts, endothelial cells, Schwann cells, MTJ/NMJ nuclei, immune, adipocytes, smooth muscle, MuSCs). For each condition, we used the RNA assay data slot (log-normalized counts), retained genes detected in ≥10 cellsand harmonized the LR database by enforcing unique Symbol rownames. CellChat objects were created followed by subsetData(), identifyOverExpressedGenes(), and identifyOverExpressedInteractions(). Communication probabilities were estimated with computeCommunProb (raw.use = TRUE, population.size = TRUE, type = “truncatedMean”), aggregated to pathways via computeCommunProbPathway() and aggregateNet(), and then compared conditions after merging objects with mergeCellChat(). We contrasted the number and overall strength of interactions using compareInteractions() and visualized rewiring with circle plots, global heatmaps, and netVisual_diffInteraction(). Sender or receiver roles were quantified with netAnalysis_computeCentrality() and netAnalysis_signalingRole_scatter(), and condition-specific changes were profiled for myonuclei using netAnalysis_signalingChanges_scatter(). Pathway architecture was evaluated by computing functional and structural similarities with computeNetSimilarityPairwise(), embedding joint manifolds via netEmbedding(), clustering with netClustering(), and summarizing distances/rankings using rankSimilarity() and rankNet(measure = “weight”). Differential LR analysis on the merged object used identifyOverExpressedGenes(thresh.pc = 0.1, thresh.fc = 0.05, thresh.p = 0.05), mapping DE results back to communications with netMappingDEG() and extracting LR sets via subsetCommunication() (ligand.logFC thresholds: +0.05 for up-regulated in KPC, −0.05 for down-regulated). Regulated LR programs were visualized with bubble plots, chord diagrams, and word clouds, and gene expression distributions were compared across groups with plotGeneExpression.

### SCENIC and transcription network analysis

Single-nucleus RNA-seq data were processed in Seurat (v5.3.0) (76). Nuclei passing basic QC (same as above) were retained, log-normalized, and highly variable genes were identified with FindVariableFeatures (vst). For TF program inference we ran pySCENIC (v0.12.1) on Linux/WSL2 in three steps: (i) GRN inference with pyscenic grn using GRNBoost2 and a curated mouse TF list (TF_symbols.txt supplied with pySCENIC resources); (ii) motif enrichment/pruning with pyscenic ctx against the mm10 cisTarget ranking databases (v10_clust; 10 kb up/10 kb down and 500 bp up/100 bp down) and the v10nr motif annotations table, combining promoter-proximal and distal evidence; and (iii) AUCell scoring with pyscenic aucell to produce per-cell regulon activity in a .loom. SCENIC outputs were imported into R and the AUCell matrix (RegulonsAUC × CellID) was added as a Seurat assay (RegAUC). Loom Cell IDs were aligned to Seurat barcodes, and basic coverage checks confirmed that TFs/targets from the GRN were present in the expression matrix.

Differential regulon activity between KPC and PBS groups was tested on RegAUC with two-sided Wilcoxon tests and Benjamini–Hochberg (BH) correction (logfc.threshold = 0 to retain subtle activity shifts; downstream filters control specificity). Unless noted, these comparisons were performed across all myofiber nuclei; cluster-stratified analyses were additionally performed for cachexia-associated clusters (e.g., clusters 2 and 14) where indicated. “Triple-evidence” TFs were defined as those that satisfied all three criteria: (i) significant up-regulation of the TF’s regulon activity in cachectic myofibers (RegAUC, BH FDR < 0.05); (ii) overlap between SCENIC-inferred high-confidence targets and a high-confidence cachexia DEG set from RNA (Wilcoxon, BH FDR < 0.05, avg_log2FC > 0; computed within clusters 2/14 for atrophy-focused contrasts); and (iii) up-regulation of the TF’s own transcript (RNA, BH FDR < 0.05). For visualization, regulon AUCs were row-wise z-scored and hierarchically clustered (ComplexHeatmap); UMAP feature maps were drawn for key regulons (RNA-based PCA/UMAP for display), and AUCell thresholds were estimated per regulon (AUCell_exploreThresholds) to binarize activity and compute the fraction of active cells by cluster/condition. TF cooperativity within cachexia-associated clusters was assessed using Pearson correlations of RegAUC followed by Ward.D2 clustering to define modules. High-confidence targets of triple-evidence TFs underwent GO Biological Process enrichment (clusterProfiler enrichGO, mouse OrgDb; BH-adjusted q ≤ 0.05). An enrichment map linked terms by Jaccard overlap ≥ 0.2, followed by Louvain community detection and calculation of degree, betweenness, and closeness centralities; terms explicitly referring to TORC1/mTOR were annotated as “core mTOR”, and translation/ribosome terms as “indirect translation/ribosome”. To connect TFs to functions, a Sankey diagram linked triple-evidence TFs to enriched GO terms with ribbon width proportional to the number of high-confidence targets contributing to each term. A directed TF-TF network was generated by treating SCENIC targets that are TFs as downstream nodes (self-loops removed) and plotted with a force-directed layout (ggraph/igraph; node size = out-degree). Unless stated otherwise, tests were two-sided with BH correction.

### Augur-based perturbation sensitivity

We quantified cell-state sensitivity to cachexia with Augur on a Seurat object (77). Cell types were defined from seurat_clusters, and conditions were KPC vs PBS (from orig.ident). For each cell type, Augur’s calculate_auc trained a random-forest classifier to distinguish KPC from PBS using repeated subsampling (50 iterations; 20 cells per class) with 3-fold cross-validation. Area-under-ROC (AUC) values were averaged across subsamples to rank perturbation sensitivity. Analyses were performed in R (Seurat v5.3.0, Augur v1.0.3).

### Pathway module scoring and visualization

We quantified pathway-level activity per nucleus using curated gene sets for autophagy, the ubiquitin–proteasome system (UPS), oxidative phosphorylation (OXPHOS) and angiogenesis. Metadata were harmonized by renaming orig.ident to KPC or PBS, and coarse cell types were derived from Seurat clusters (e.g., Type I, Type IIa/IIb/IIx, cachectic myonuclei). For each gene set, per-cell module scores were computed with Seurat AddModuleScore on the RNA assay (78). Gene-level expression was summarized with DotPlot grouped by cell type and split by condition (KPC vs PBS). Pathway activity distributions were visualized with VlnPlot for the corresponding module score (split by condition across cell types).

### Statistical Analysis

All wet-lab data are presented as mean ± standard error of the mean (SEM). Statistical analyses were performed using GraphPad Prism 10.0. Comparisons between two groups were conducted using two-tailed unpaired Student’s t test. For experiments involving multiple comparisons, two-way analysis of variance (ANOVA) followed by Tukey’s post hoc test was applied. *p* ≤ 0.05 was considered statistically significant. Additional statistical details, including exact n, are provided in the corresponding figure legends.

## Acknowledgements

This work was supported by the National Institute of Health grant AR081487 and CA294365 to AK.

## Authors’ contributions

AK designed the work. BX, ASJ, MTS, SL, and PTH performed the experiments and analyzed the results. BX and ASJ wrote the first draft of the manuscript. AK and other authors edited and finalized the manuscript.

