## Supplementary material for "Fiber-type vulnerability and proteostasis reprogramming in skeletal muscle during pancreatic cancer cachexia": Figures S1-10, Tables S1 and S2

**By**

**Bowen Xu, Aniket S. Joshi, Meiricris Tomaz da Silva, Silin Liu, and Ashok Kumar**

**This file contains Figure S1-S10 and Tables S1 and S2.**

#### Supplemental Figures' Legends.

##### FIGURE S1

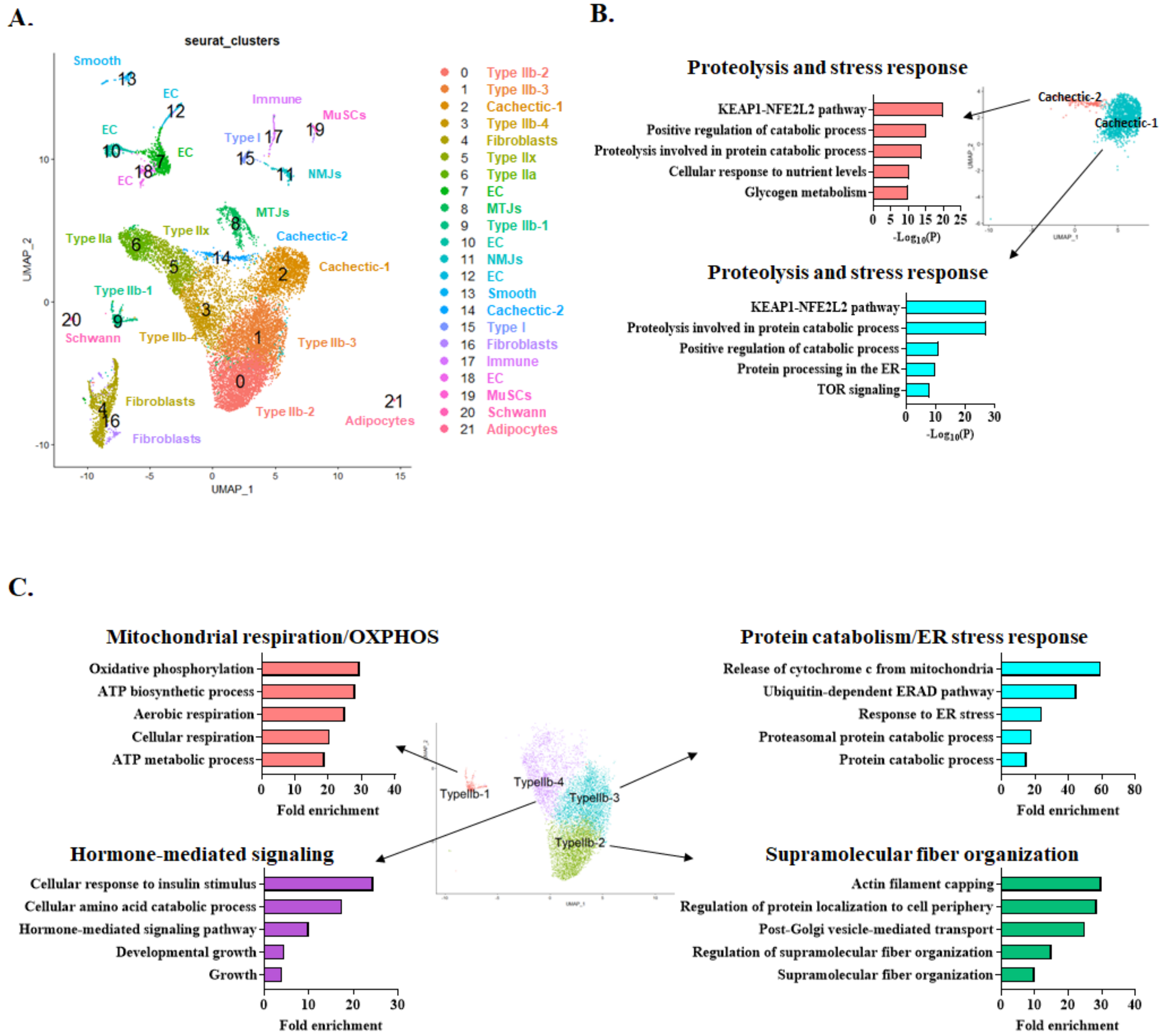


**Figure S1. Annotation of nuclear clusters to different cell types in skeletal muscle. (A)** UMAP visualization of the integrated single-nucleus RNA-seq dataset from skeletal muscle of control and KPC tumor-bearing mice following unsupervised clustering. A total of 21 transcriptionally distinct clusters were resolved, each point representing a single nucleus, with colors indicating cluster identity. **(B)** Enrichment analysis of the two cachectic clusters to annotate their differences. **(C)** Enrichment analysis of the four type-IIb clusters to annotate their differences.

**FIGURE S2**

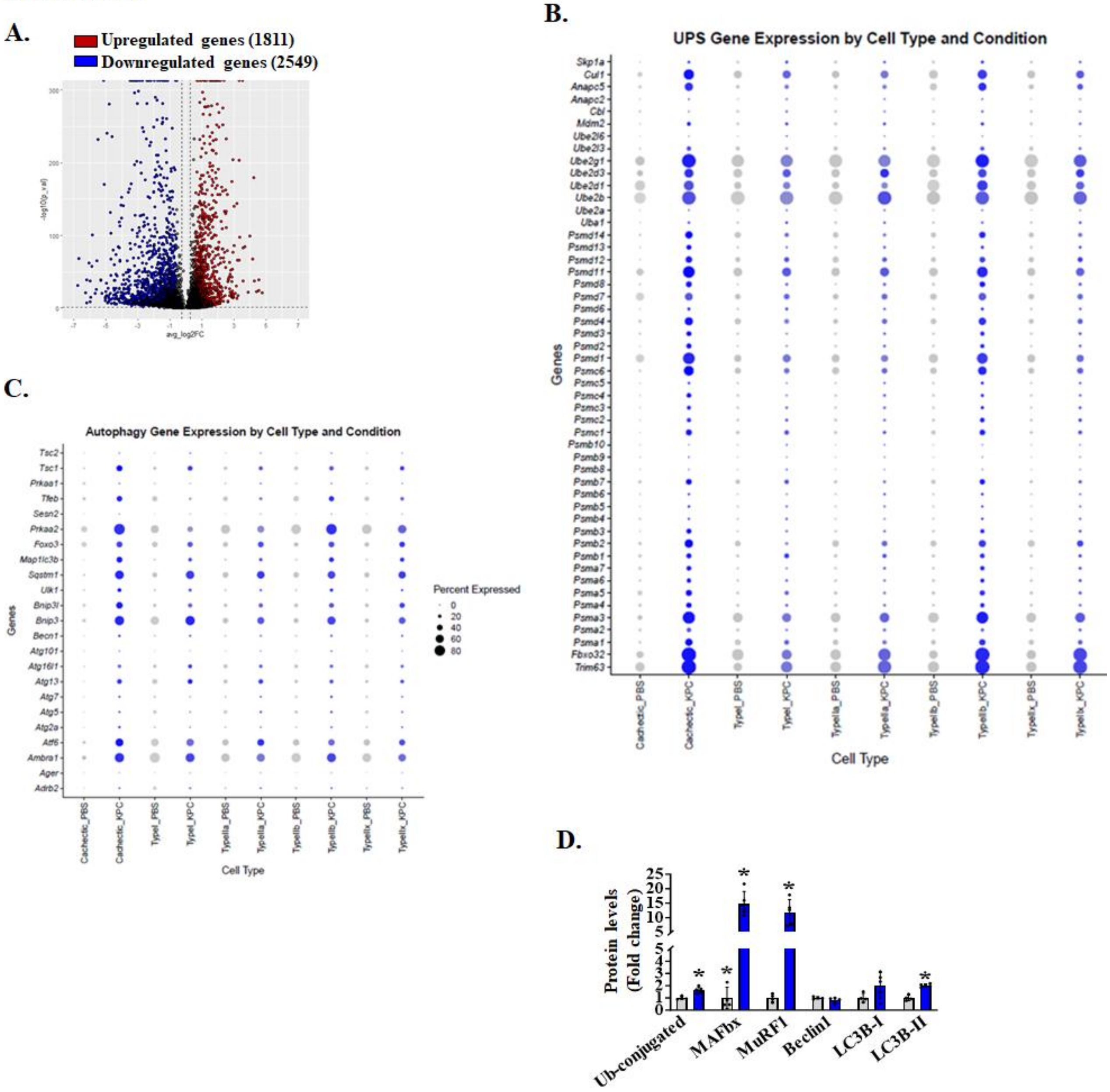


**Figure S2**. **Increased expression of UPS and autophagy-related genes in cachectic myonuclear clusters. (A)** Volcano plot showing differentially expressed genes in cachectic myonuclei compared with all control muscle nuclei, with 1,811 genes upregulated (red) and 2,549 genes downregulated (blue). **(B)** Dot plot showing the expression patterns of genes within the ubiquitin–proteasome system (UPS) gene set used for module scoring. **(C)** Dot plot showing the expression patterns of genes within the autophagy gene set used for module scoring. Dot size indicates the percentage of nuclei expressing each gene within a given fiber type, and dot color reflects average expression levels. **(D)** Quantification of the levels of total ubiquitinated proteins, MAFbx, MuRF1, Beclin1, LC3B-I/II.

**FIGURE S3**

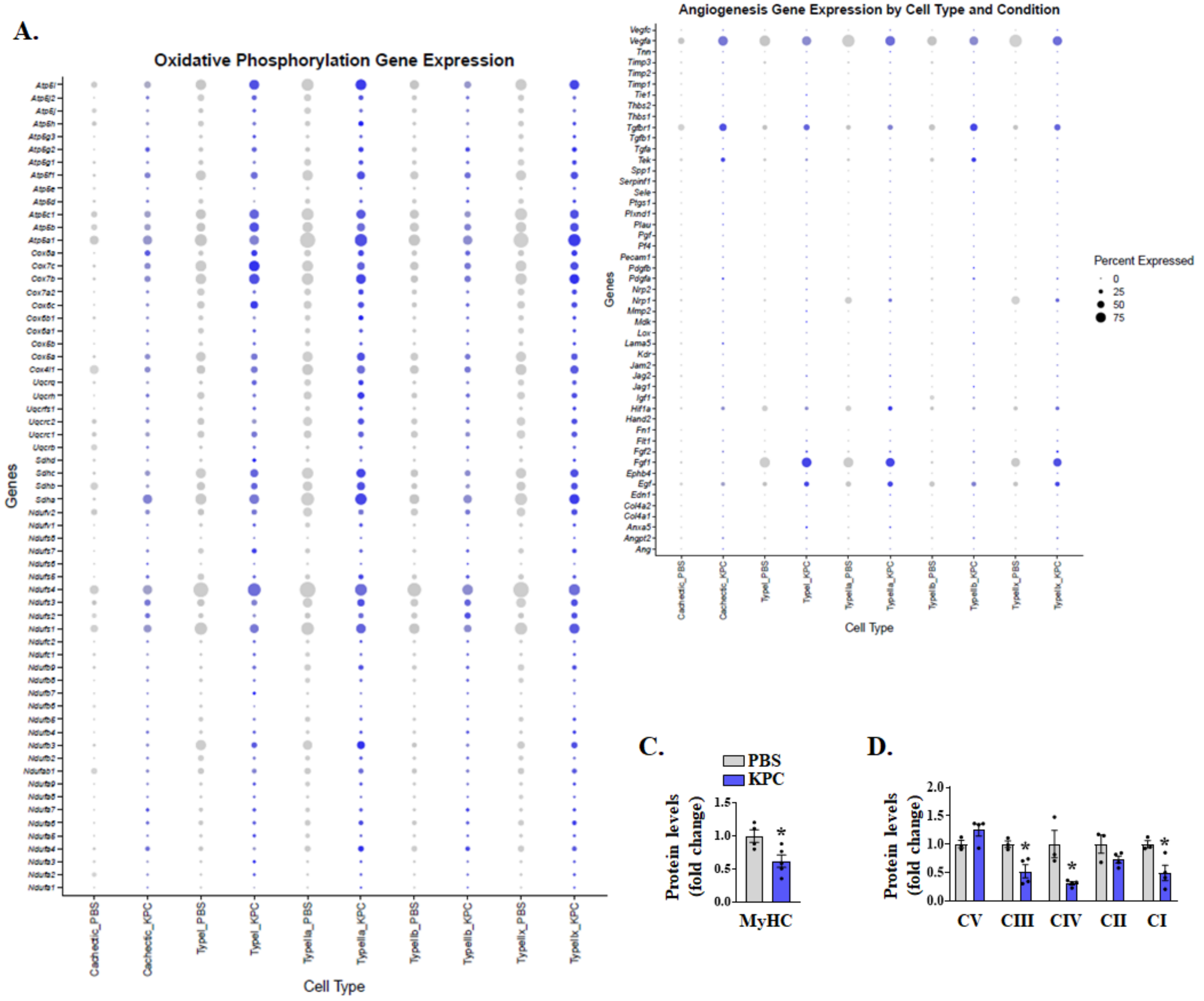


**Figure S3. Repression in gene expression of molecules related to oxidative phosphorylation and angiogenesis.** **(A)** Dot plot showing the expression patterns of genes within the oxidative phosphorylation gene set used for module scoring. **(B)** Dot plot showing the expression patterns of genes within the angiogenesis gene set used for module scoring. Dot size indicates the percentage of nuclei expressing each gene within a given fiber type, and dot color reflects average expression levels. Relative protein levels of **(C)** MyHC, and **(D)** mitochondrial oxidative phosphorylation (OXPHOS) complexes I-V.

**FIGURE S4**

**A.**

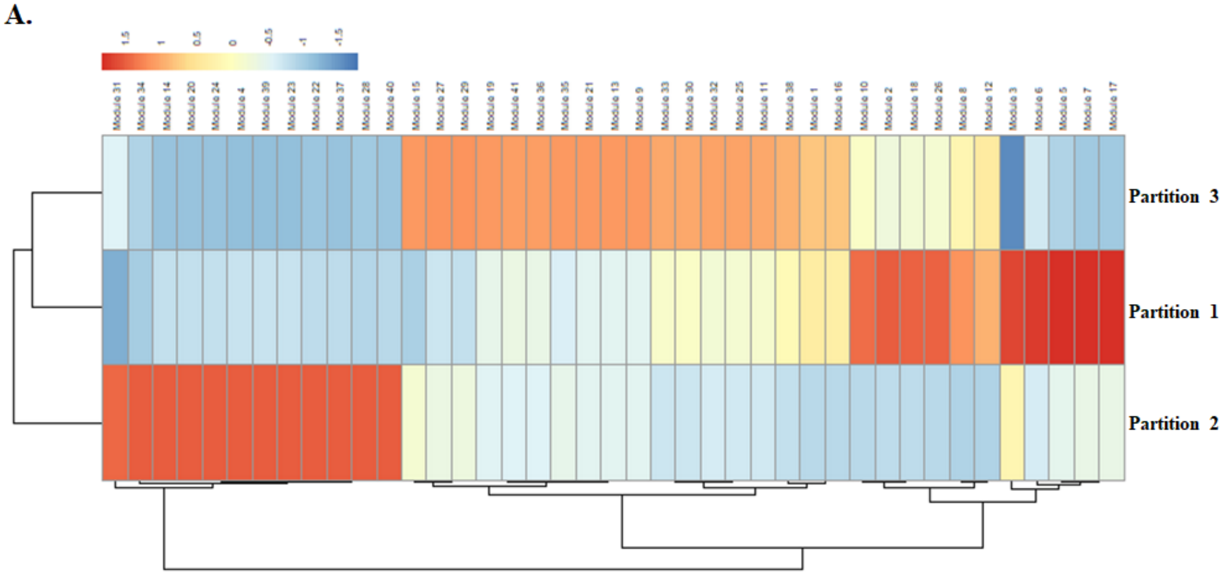


**Figure S4. Gene co-expression modules of cachectic myonuclei.** Heatmap showing 41 gene modules identified by Louvain community detection method grouped into 3 partitions in myonuclear clusters of KPC tumor-bearing mice.

### Figure S5

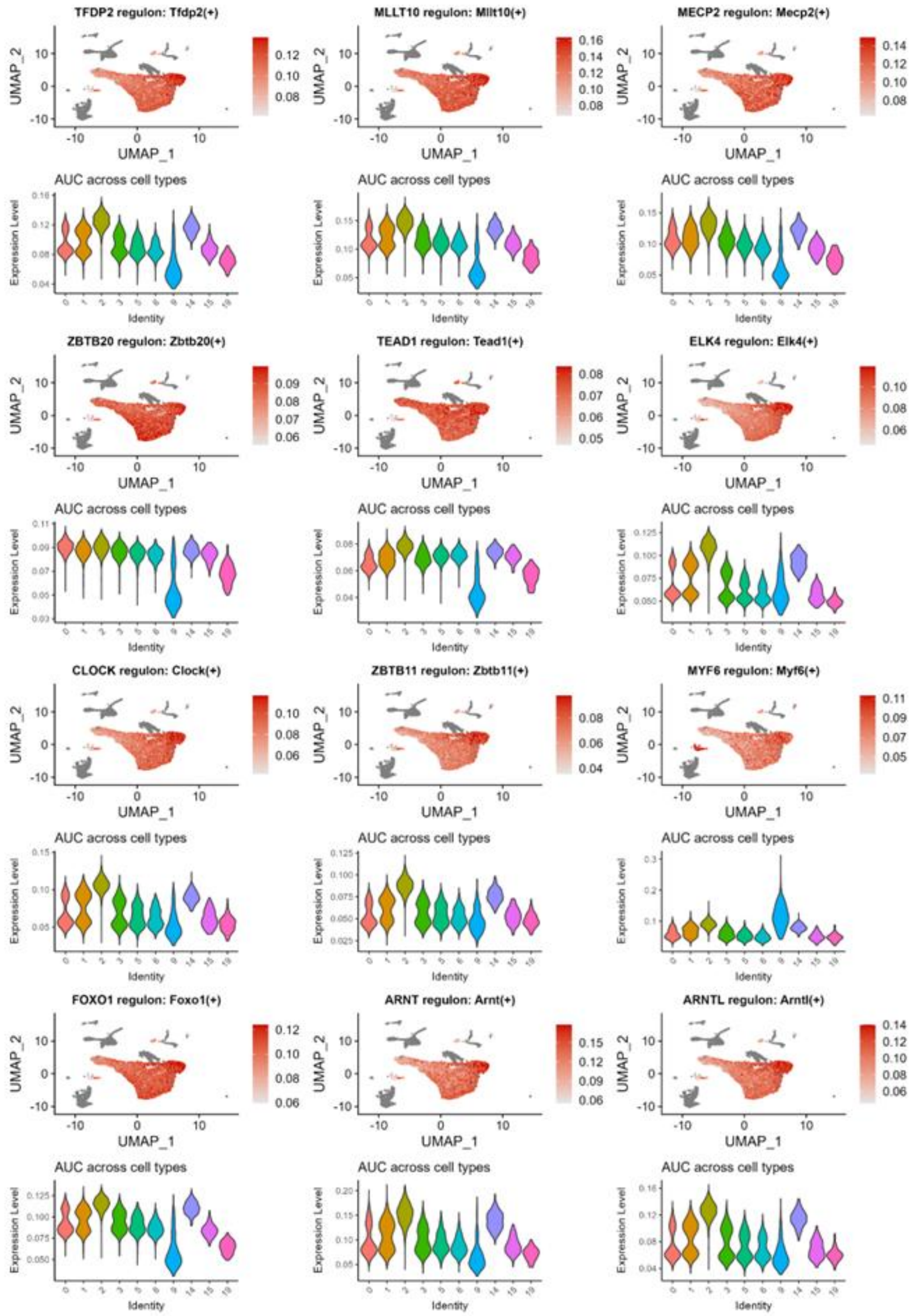


**Figure S5. Translation initiation and mTOR-related transcription factors activity in myonuclei.** UMAP visualization of regulon activity for mTOR- and translation-associated transcription factors, with violin plots showing cluster-specific activation patterns.

### Figure S6

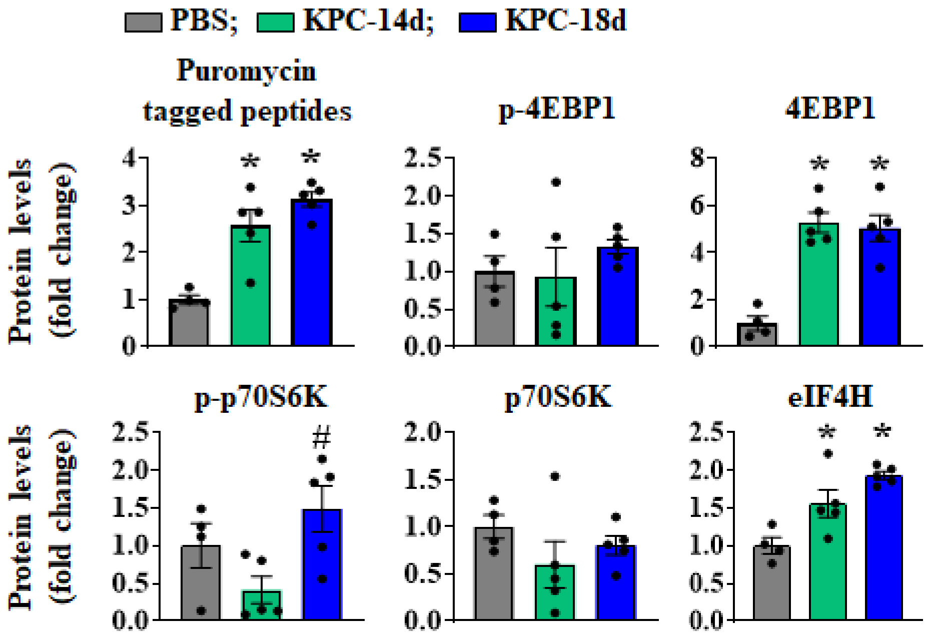


**Figure S6. KPC tumor growth induces alterations in muscle protein synthesis.** Quantification of levels of puromycin-tagged peptides and levels of p-4EBP1, 4EBP1, p-70S6K, p70S6K, eIF4H, and GAPDH in TA muscle of control and KPC tumor-bearing mice after 14 and 18 days of KPC cells injection in pancreas.

### Figure S7

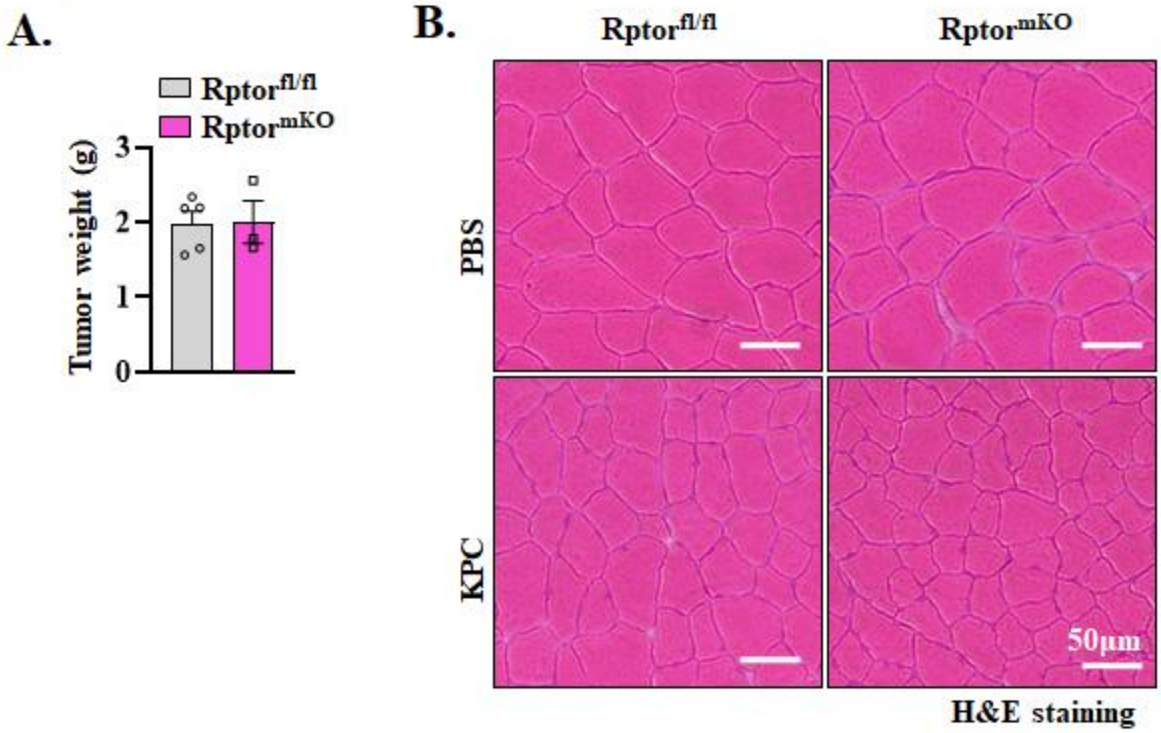


**Figure S7. Targeted deletion of Raptor exacerbates muscle loss in response to KPC tumor growth. (A)** Wet weight of tumor in $Rptor^{fl/fl}$ and $Rptor^{mKO}$ mice. **(B)** Representative images of TA muscle cross-sections after H&E staining. Scale bar, 50 μm.

#### Figure S8

A.

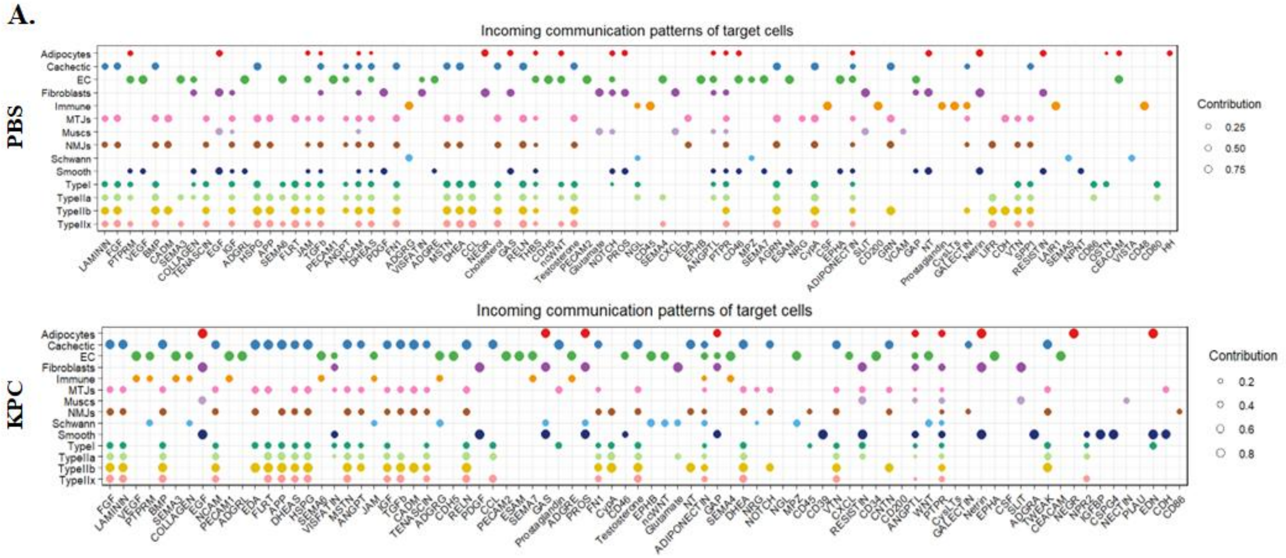


B.

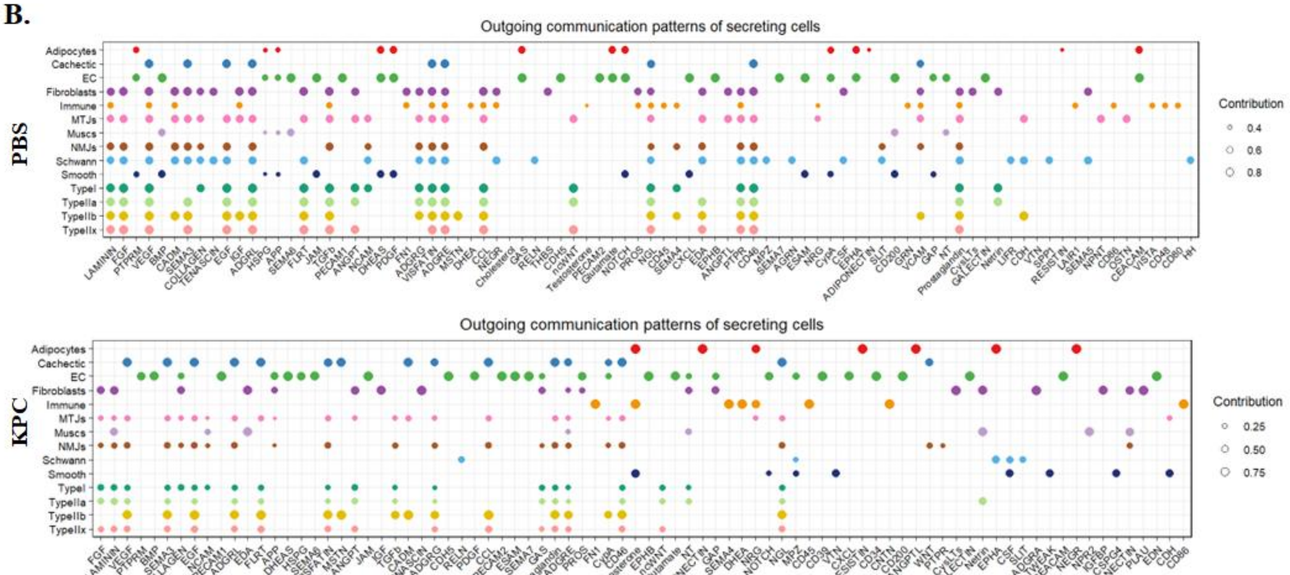


**Figure S8. Changes in intercellular communications within skeletal muscle of tumor-bearing mice. (A)** Dot plots showing incoming signaling patterns across all cell types in PBS control and KPC tumor-bearing muscle. **(B)** Dot plots showing outgoing signaling patterns across all cell types in PBS control and KPC tumor-bearing muscle.

**Figure S9**

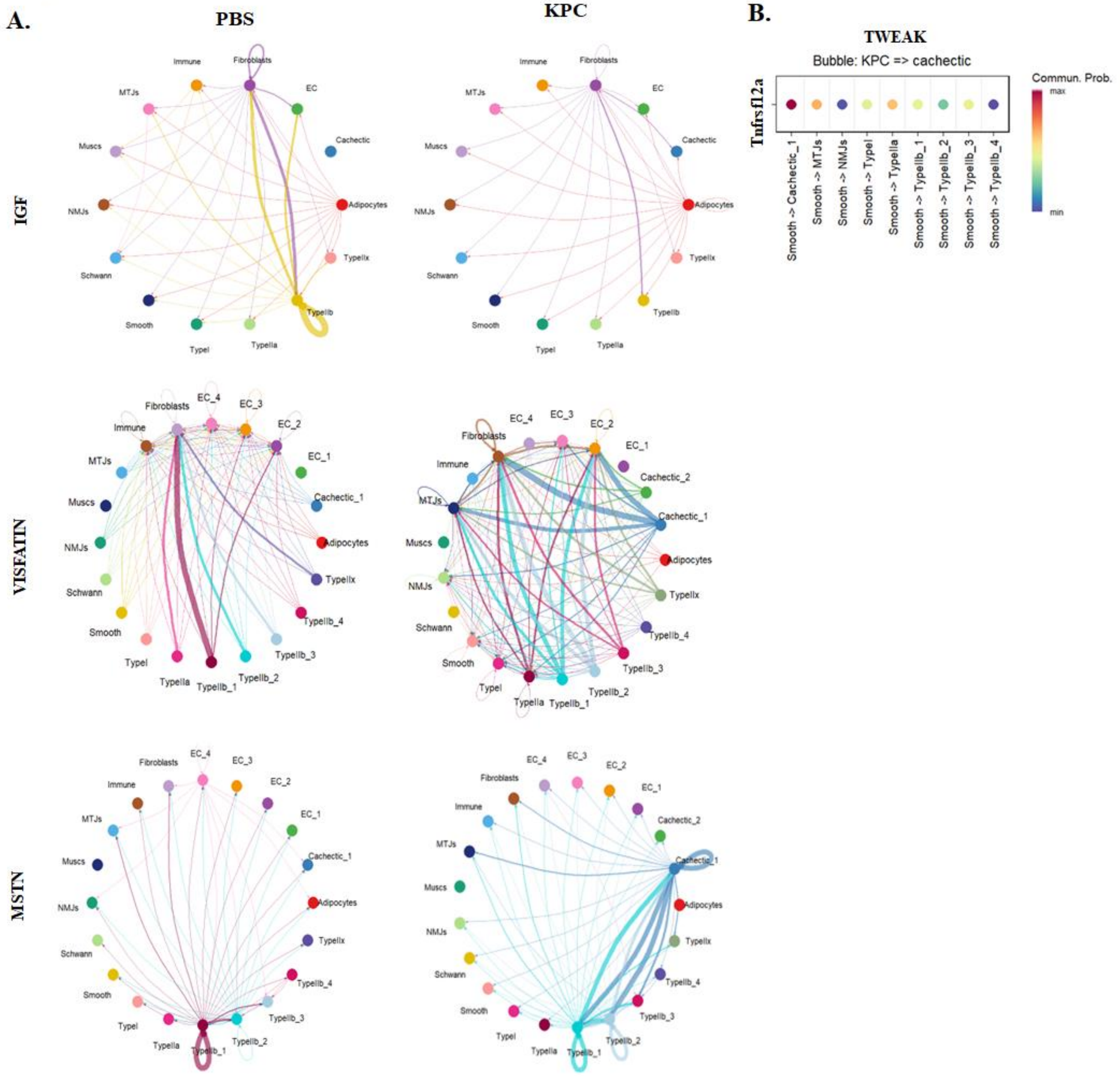


**Figure S9. Intercellular signaling changes in cachectic muscle. (A)** Circle plot illustrating intercellular communication mediated by IGF, MSTN, VISFATIN in control (PBS) and KPC tumor-bearing mice. **(B)** Dot plot showing interaction of TWEAK with its receptor Tnfrsf12a in different myonuclei of KPC tumor-bearing mice identified by CellChat analysis of snRNA-Seq dataset.

### Figure S10

#### Fig. 2F

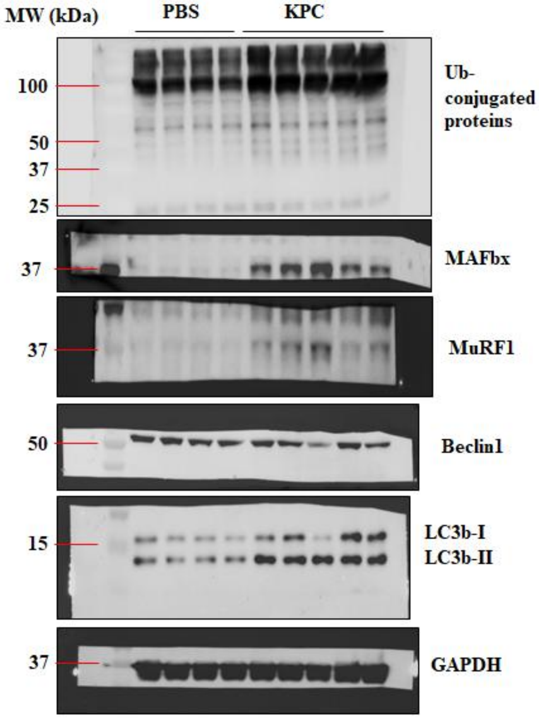


#### Fig. 3E

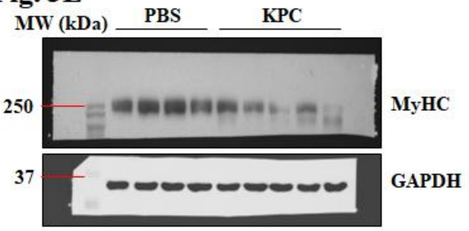


#### Fig. 6D

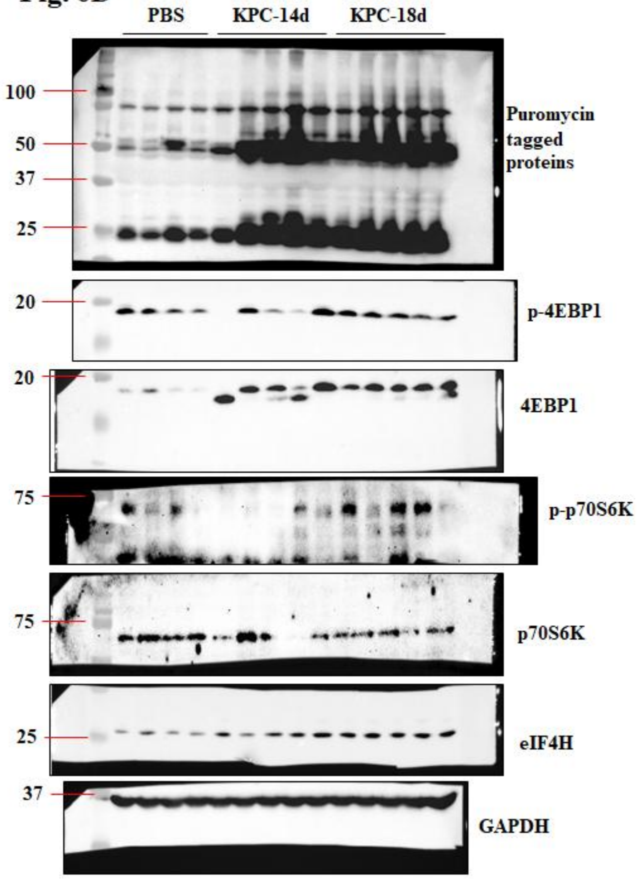


#### Fig. 3F

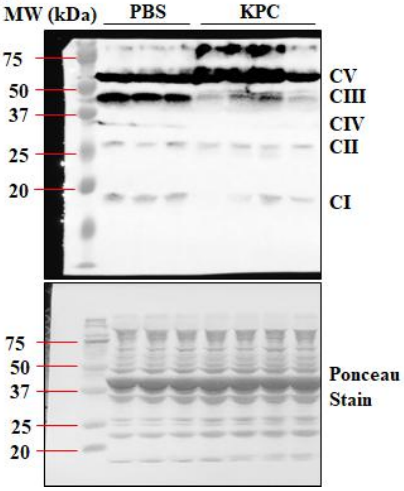

#### Figure S10 (Continuation)

##### Fig. 7L

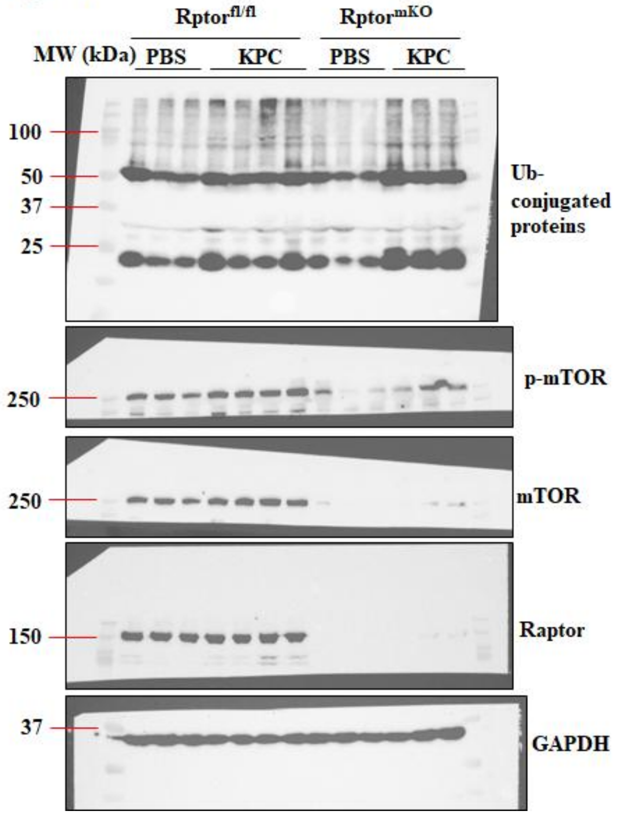


**Figure S10. Uncropped western blot gel images.**

**Table S1.** List of antibodies used for Western blot and Immunofluorescence.

| Antibody | Source and Catalog no. | Analysis |
|---|---|---|
| Anti-MAFbx | ECM Biosciences #AP2041 | WB |
| Anti-MuRF1 | R&D Systems #AF5366 | WB |
| Anti-Beclin1 | Cell Signaling #3495S | WB |
| Anti-LC3b | Cell Signaling #2775S | WB |
| Anti-GAPDH | Cell Signaling #5174S | WB |
| Anti-Type I MyHC | DSHB #BA-D5 | IF |
| Anti-Type IIa MyHC | DSHB #SC-71 | IF |
| Anti-Type IIb MyHC | DSHB #BF-F3 | IF |
| Anti-mTOR | Cell Signaling #2972 | WB |
| Anti-p-mTOR | Cell Signaling #2971S | WB |
| Anti-Raptor | Cell Signaling #2280 | WB |
| Anti-Puromycin | Millipore Sigma #MABE343 | WB |
| Anti-4EBP1 | Cell Signaling #9452S | WB |
| Anti-p-4EBP1 | Cell Signaling #2972 | WB |
| Anti-p70S6K | Cell Signaling #9202S | WB |
| Anti-p-p70S6K | Cell Signaling #9208S | WB |
| Anti-eIF4H | Cell Signaling #3469T | WB |
| Anti-Laminin | Sigma #L9393 | IF |
| Anti-rabbit IgG | Cell Signaling #7074S | WB |
| Anti-mouse IgG | Cell Signaling #7076S | WB |
| Anti-goat IgG | Invitrogen #A15999 | WB |
| Anti-mouse IgG1 AF568 | Invitrogen #A21124 | IF |
| Anti-rabbit IgG AF488 | Invitrogen #A11034 | IF |
| Goat anti-Mouse IgG2b AF350 | Life Technologies #A21140 | IF |
| Goat anti-Mouse IgG1 AF568 | Life Technologies #A21124 | IF |
| Goat anti-Mouse IgM AF488 | Life Technologies #A21042 | IF |

**Table S2.** List of primers used for PCR/qRT-PCR analysis

| Name | Forward primer (5′-3′) | Reverse primer (5′-3′) |
|---|---|---|
| β-actin | CAGGCATTGCTGACAGGATG | TGCTGATCCACATCTGCTGG |
| 18s | CGGCTACCACATCCAAGGAA | GCTGGAATTACCGCGGCT |
| 28s | TCATCAGACCCCAGAAAAGG | GATTCGGCAGGTGAGTTGTT |
| TIF1a | ATTCCCGTTTGTGAGGAAGTCCGA | TATCCTGCCGCGATACACTCACAT |
| PAF53 | TCAGAACAAGACTTTCAGGGACAA | CTGCTTGGTGCTTCCAAAGG |
| Polr1b | TGGGAATCTGCGTTCTAAAAC | TTCAGCTTGTCAGCCACAACA |
| UBF | CGCGCAGCATACAAAGAATAC | GTTTGGGCCTCGGAGCTT |
| Rpl5 | GGAGGTGAATGGAGGTGAATA | AGTTGTAGTTCGGGCAAGAC |
| Rpl11 | TGCCTCAATATCTGCGTCGG | TTCTCCGGATGCCAAAGGAC |
| Rps3 | CAGAGAAAGTGGCCACAAGA | TGAACCGAAGCACACCATAG |
| Rps6 | CATGGGGGAGTTGGACCATA | TGGCCCTCTGTTCACACTA |
| Arrdc3 | GCATAAAGGGCATTTGGTGGT | ATGGTAGTGAGTGCCCAAGG |
| Sik1 | CCAAACACCTCTGGAAGGAA | CCTTTCCTCTCCCTTAATGACC |
| Sesn1 | CTATTTGGCTGAGTGCTGCTA | GTCTTGACTGAGCTGTGAGTAA |
| Retreg1 | CCATGAGTGATGTGGTCATTGG | AGACTGCAGTGGCAGAGTAT |
| Egln3 | GAATCCCACAAGCAGCATAGA | GACAGTTCTTCCTGGACACTTT |
| Acss1 | GCTGTGATCTGGAAGGAGTTT | ATGGGACCAGAGGCTTAGTA |
| Rgcc | CGAAGACTTCATTGCCGATCT | TCTCCCACAGCAGATTCTAGT |
| Irs2 | TTGCTGTGTAGCTGAATCCC | ATATGCCATCTGTGTCCTGTG |
